# Developmentally regulated alternate 3’ end cleavage of nascent transcripts controls dynamic changes in protein expression in an adult stem cell lineage

**DOI:** 10.1101/2022.05.09.489277

**Authors:** Cameron W. Berry, Gonzalo H. Olivares, Lorenzo Gallicchio, Gokul Ramaswami, Alvaro Glavic, Patricio Olguín, Jin Billy Li, Margaret T. Fuller

## Abstract

Alternative polyadenylation (APA) generates transcript isoforms that differ in the position of the 3’ cleavage site, resulting in the production of mRNA isoforms with different length 3’UTRs. Although widespread, the role of APA in the biology of cells, tissues and organisms has been controversial. We identified over 500 *Drosophila* genes that express mRNA isoforms with a long 3’UTR in proliferating spermatogonia but a short 3’UTR in differentiating spermatocytes due to APA. We show that the stage-specific choice of the 3’ end cleavage site can be regulated by the arrangement of a canonical polyadenylation signal (PAS) near the distal cleavage site but a variant or no recognizable PAS near the proximal cleavage site. The emergence of transcripts with shorter 3’UTRs in differentiating cells correlated with changes in expression of the encoded proteins, either from off in spermatogonia to on in spermatocytes or vice versa. Polysome gradient fractionation revealed over 250 genes where the long 3’UTR versus short 3’UTR mRNA isoforms migrated differently, consistent with dramatic stage-specific changes in translation state. Thus, the developmentally regulated choice of an alternative site at which to make the 3’end cut that terminates nascent transcripts can profoundly affect the suite of proteins expressed as cells advance through sequential steps in a differentiation lineage.

## Introduction

The switch from proliferation to differentiation is a key event in both development and adult tissue renewal. Failure to cleanly shut down proliferation programs may contribute to the initiation of oncogenesis in the adult stem cell lineages that maintain short-lived differentiated cell populations and/or repair many tissues in the body. Conversely, delay or defects in turning on proper genetic programs for cell type-specific differentiation may lead to tissue dysmorphosis, degenerative disease and aging.

Alternative mRNA processing resulting in cell type-specific mRNA isoforms may contribute to changes in cell state during differentiation (Ji et al. 2009; Di Giammartino et al. 2011; Lutz and Moreira 2011; Elkon et al. 2013; Gruber and Zavolan 2019; Cheng et al. 2020; Agarwal et al. 2021; Pereira-Castro and Moreira 2021; Sommerkamp et al. 2021). In particular, the selection of alternative sites at which to make the 3’end cut that terminates the nascent transcript (termed alternative polyadenylation – APA) leads to expression of mRNA isoforms with different 3’UTR lengths (Di Giammartino et al. 2011; Shi 2012; Mueller et al. 2013). APA has been associated with specific cell types or disease states (Ji et al. 2009; Ji and Tian 2009; Mayr and Bartel 2009; Elkon et al. 2012; Morris et al. 2012; Gruber and Zavolan 2019; Mohanan et al. 2021).

For example, developmentally regulated APA at several genes leads to the expression of transcript isoforms with much longer 3’UTRs in the brain than in other tissues (Ji et al. 2009; Hilgers et al. 2011; Derti et al. 2012; Smibert et al. 2012; Bae and Miura 2020), while oncogenic transformation has been correlated with emergence of transcript isoforms with short 3’UTRs due to APA (Mayr and Bartel 2009; Fu et al. 2011).

Although changes in 3’UTR length due to APA have been widely reported, including in *Drosophila* testes (Smibert et al. 2012), their biological relevance is poorly understood and has been extensively debated (Xu and Zhang 2020). While 3’UTRs can harbor *cis*-regulatory information that controls translation of the mRNA, the extent to which widespread alternative 3’ end cleavage alters the proteome has been investigated in only a limited number of biological systems. For a small number of specific genes, transcript isoforms produced by alternative 3’ end cleavage are differentially translated in human cancer cell lines (Mayr and Bartel 2009). Two studies, one of global translation in the mouse NIH 3T3 cell line (Spies et al. 2013) and the other of global protein abundance during the activation of murine T-cells (Gruber et al. 2014), both indicated that alternative 3’ end cleavage leading to different length 3’UTRs had relatively little contribution to differential mRNA translation or protein abundance in the cell types assessed. On the other hand, a study of global translation in five human and two mouse cell lines found that transcript isoforms processed with short 3′ UTRs show higher translational efficiency than those with long 3′ UTRs in all tested lines except for NIH3T3 cells (Fu et al. 2018). Another study analyzing ribosomal association of transcript isoforms in human HEK 293T cells found that mRNA isoforms with long 3’UTRs were associated with lower protein synthesis (Floor and Doudna 2016) than short 3’UTR isoforms from the same gene. Thus, over a decade after the identification of widespread APA, it has remained unclear whether biological systems employ developmentally regulated alternative 3’ end cleavage as a mechanism to specify large-scale changes in protein expression in different cell types.

Here, using the *Drosophila* male germline as a model adult stem cell lineage (Figure 1A), we show that developmentally regulated alternative 3’ end cleavage leading to the production of transcript isoforms with shortened 3’UTRs alters the translation state of many mRNAs in differentiating cells compared to their proliferating precursors. We found that over 500 genes produce mRNA isoforms with long 3’UTRs in proliferating spermatogonia but short 3’UTRs soon after initiation of the program for meiosis and gamete differentiation in spermatocytes. Strikingly, differences in the behavior of mRNA isoforms with short vs long 3’UTRs in polysome fractionation studies suggested that for at least half of the genes, 3’UTR shortening due to APA is accompanied by a change in translation state. For 50 genes, the long 3’UTR isoform expressed in testes enriched for proliferating spermatogonia co-migrated with sub-ribosomal light fractions, suggesting the mRNA was not being translated, while the partner isoform with truncated 3’UTR expressed in testes enriched for differentiating spermatocytes co-migrated with mono-, di-, or polysomes. For another 200 genes, transcript isoforms with a long 3’UTR expressed in spermatogonia co-migrated with mono-, di-, or polysomes, while their partner short 3’UTR mRNA isoforms produced from the same gene in young spermatocytes migrated in lighter, sub-ribosomal fractions, suggesting lack of translation. Of these 200 genes, for 139 the short 3’UTR isoform moved from lighter, sub-ribosomal fractions in testis extracts enriched for young spermatocytes to co-migration with 80S or higher fractions in extracts enriched for maturing spermatocytes. This stage-specific ON -> OFF -> ON co-migration of mRNA isoforms with ribosomes suggests a rationale for why the APA mechanism may be especially useful for dynamic regulation of protein production in a differentiation sequence: while 3’UTR shortening by APA can specify a sharp decrease in protein production in early spermatocytes, transcription continues, providing mRNAs that can be translated at later stages, such as maturing spermatocytes or spermatids undergoing morphogenesis. Our results suggest that developmentally regulated APA may provide a mechanism to facilitate clean and rapid transitions between developmental states via dynamic translational regulation, turning off the production of specific proteins for the prior program that may be deleterious for early steps of differentiation, while maintaining transcripts and so the ability to re-activate translation at later stages of differentiation.

**Figure 1.**
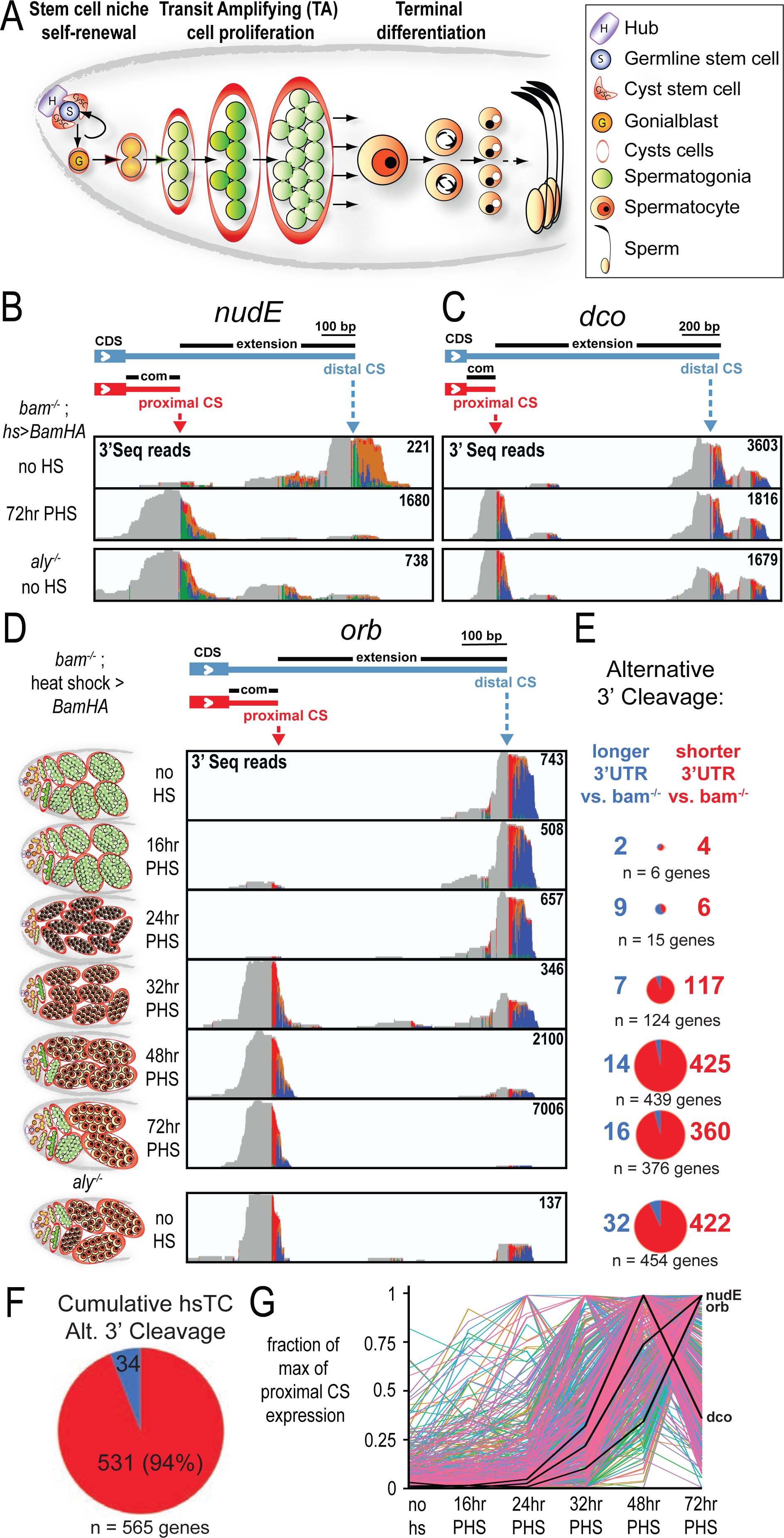
Differential 3’ end cleavage of selected transcripts is associated with the switch from spermatogonia proliferation to spermatocyte differentiation in *Drosophila*. (A) Main stages of *Drosophila* male germ cell differentiation. (B-C) 3’ Seq tracks from one of two biological replicates plotted on the 3’ genomic region of (B) *nudE* and (C) *dco* from testes from *bam ^Δ86^*^/1^; *hs-Bam-HA* (*bam* heat shock time course (hsTC)) flies with no heat shock or 72 hours post heat shock (PHS), as well as testes from *aly* mutant flies with no heat shock. The maximum number of supporting reads are indicated on the right of each track. On top, a gene model representing the two most abundant isoforms of 3’UTR for each gene, indicating the position of the proximal (red) and distal (blue) cleavage site (CS) according to peaks from 3’ Seq tracks. (D) 3’ Seq tracks of one of two biological replicates plotted on the 3’ genomic region of *orb* from the *bam* hsTC flies at indicated times PHS and from *aly* mutant testes. On the left, testes diagrams indicating the cell diversity and developmental stage at each timepoint. (E) Pie charts (size normalized to gene list) for each timepoint indicating the number of genes detected as undergoing alternative 3’ cleavage events resulting in longer (blue) or shorter (red) 3’UTRs relative to in testes from *bam ^Δ86^*^/1^; *hs-Bam-HA* flies without HS. Bottom row: comparison of *aly^5p/2^* flies without heat shock to *bam^Δ86^*^/1^*; hs-Bam-HA* flies without HS. (F) Cumulative pie chart containing all genes detected as changing 3’UTR cut site in the *bam* hsTC. (G) Line graph of the 531 genes called as showing stage-specific alternative 3’ cleavage, with the relative level of the short 3’UTR isoform plotted as the fraction of maximum value over the time course. Black lines: relative short 3’UTR transcript levels for *nudE*, *orb*, and *dco*.

## Results

### Stage-specific 3’UTR shortening by APA in the *Drosophila* male germline stem cell lineage

Sperm are produced in large quantities throughout reproductive life in a robustly active adult stem cell lineage. The most dramatic changes in gene expression in this lineage occur when proliferating spermatogonia complete a final mitosis, undergo a last (premeiotic) S-phase and initiate the cell growth and differentiation program characteristic of spermatocytes. In *Drosophila* (Figure 1A), male germline stem cells located at the apical tip of the testes normally divide one at a time, producing a replacement stem cell and a gonialblast, which becomes enclosed in a pair of somatic cyst cells and initiates spermatogonial proliferation. After four rounds of synchronous transit-amplifying mitotic divisions, the resulting 16 interconnected germ cells undergo the premeiotic S-phase in synchrony and then embark on the 3.5-day spermatocyte program, which takes place during the meiotic prophase. The spermatocytes grow 25X in volume and express many transcripts required for the meiotic divisions and the extensive cellular morphogenesis that takes place in the resulting haploid spermatids (Fuller 1993). Wild-type testes contain a continuous stream of differentiating germ cells, each germline cyst at a different stage, making molecular analysis of stage-specific events challenging. However, germ cells can be induced to switch from proliferating spermatogonia to the onset of spermatocyte differentiation in meta-synchrony *in vivo* using a *bam^-/-^;hs-Bam* time course system (Kim et al. 2017).

Briefly, in males mutant for *bag-of-marbles* (*bam*), testes fill with spermatogonial cysts in which the germ cells continue to proliferate and eventually die, never becoming spermatocytes. To induce semi-synchronous differentiation of spermatogonia to spermatocytes *in vivo*, *bam* mutant flies carrying a heat-shock inducible transgene driving expression of Bam were subjected to a single, 30-minute pulse of heat shock at 37°C, then returned to 25°C and followed over time. Under this experimental setup, the spermatogonia complete their final mitoses and pre-meiotic DNA replication by 24 hours post heat shock (PHS), start to express markers of the early spermatocyte transcription program by 32hr PHS, are filled with early, polar spermatocytes by 48hr PHS, and with mid-stage maturing spermatocytes by 72hr PHS. The wave of differentiating germ cells enters the first meiotic division by 102hr PHS and ultimately differentiates into functional sperm by 12 days PHS (Kim et al. 2017). Note that because the *bam^-/-^;hs-Bam* flies are returned to 25°C after the initial heat shock, *bam^-/-^*spermatogonia begin to accumulate again in the testes, so that testes from males 48hr and even more so at 72hr PHS contain both differentiating spermatocytes and newly formed spermatogonia (see Figure 1D, left side diagrams).

To globally assess the changes in mRNA isoforms expressed due to APA as germ cells progress from spermatogonial proliferation to onset of the differentiation program in spermatocytes, we mapped the 3’ end cleavage sites of mRNAs expressed at different stages in male germ cell differentiation using 3’Seq, a modified version of RNA-Seq that allows precise mapping of 3’ cut sites (Beck et al. 2010). Briefly, cDNAs representing short mRNA fragments including the polyA tail were sequenced in the sense direction and transcript 3’ ends identified by plotting sequence reads where the upstream part of the read matched a unique site in the genome, but the downstream part contained multiple A residues not encoded in the genome (Figures 1B, C and D). The analysis pipeline employed to map 3’ cleavage sites required mapped reads to contain a stretch of at least 15 contiguous A residues with at least 3 not matching the genome (Figure S1A, B and Materials and Methods). To study alternative 3’ cleavage events that alter the length of 3’UTRs rather than the protein-coding sequence, we focused on 3’Seq reads that mapped to 3’UTR regions as defined by Flybase version r6.36 plus up to 500 bp downstream (Figure S1J). Using these criteria, the number of 3’ end cleavage sites called per transcript was most often 1 (Figure S1K).

Comparing the 3’ends of mRNAs expressed in testes from *bam^-/-^;hs-Bam* flies before heat shock versus 48 or 72hr PHS revealed a set of ∼500 genes that expressed mRNA isoforms with long 3’UTRs at the 0hr time point when the testes are filled with proliferating spermatogonia, but novel mRNA isoforms with shorter 3’UTRs at 48hr or 72hr PHS, when the testes have many early or mid-stage spermatocytes in addition to spermatogonia. For example, plotting the 3’Seq reads on genomic regions starting just before the stop codon and extending downstream for *nudE* (ortholog of mammalian NDEL1 (NudE Neurodevelopment Protein 1 Like 1) (Sasaki et al. 2000; Wainman et al. 2009)) and *discs overgrown (dco)* (ortholog of mammalian CKIε (Jursnich et al. 1990; Kloss et al. 1998)) (Figure 1B, C) revealed that the 3’end cut site utilized in testes filled with spermatogonia mapped 615nt (*nudE*) or 1,557nt (*dco*) downstream of the stop codon. In contrast, 3’Seq from testes 72hr PHS featured a new 3’end cut and polyadenylation site much closer to the stop codon (121nt for *nudE*; 122nt for *dco*). Thus the alternative 3’UTR isoforms shared a short common region from the stop codon to the proximal cleavage site detected at 72hr PHS, but the main transcript expressed in 0hr *bam^-/-^;hs-Bam* testes had a substantially longer 3’UTR extending to a more distal cleavage site (Figure 1B, C). The 3’UTR shortening by APA was not due to the heat shock treatment, as similar short 3’UTR mRNA isoforms were also observed in testis from flies that had not been subjected to heat shock but were mutant for the tMAC component *aly*, so the testes were filled with mature spermatocytes (Figure 1B, C). For three selected genes tested, analysis by qRT-PCR of transcript isoforms expressed in mutant backgrounds that cause accumulation of spermatogonia or accumulation of arrested late spermatocytes validated the changes in 3’UTR length during differentiation from spermatogonia to spermatocytes indicated by the 3’Seq results in the time course (Figure S2).

To determine if most of the alternative polyadenylation (APA) events leading to the expression of mRNA isoforms with shortened 3’UTRs occurred at the same time in spermatocyte differentiation, we carried out 3’Seq analysis of testes from *bam^-/-^;hs-Bam* flies at 16hr, 24hr, 32h, 48hr, and 72hr PHS. Strikingly, for most genes subject to 3’UTR APA, the switch to an alternative 3’ end cut site (i.e., proximal CS) occurred early in spermatocyte differentiation, in most cases by 48h PHS, when the testes are filled with polar spermatocytes (Figure 1D, E). Genome-wide 3’Seq analysis detected only 6 genes that had switched to an alternative 3’end cut site by 16hr PHS and 15 genes that had switched by 24hr, based on our analysis criteria (Materials and Methods). However, comparing 3’Seq data from 32hr PHS, the time point at which many spermatocytes specific mRNA markers were first detected, to *bam^-/-^;hs-Bam* testes that were filled with proliferating spermatogonia because they had not been subjected to heat shock identified 124 genes that undergo alternative 3’ cleavage, with 117 (94.4%) producing transcripts with shorter 3’UTRs at 32h PHS (Figure 1E). By 48hr PHS, when the testes were filled with young spermatocytes at the polar spermatocyte stage, a peak number of 439 genes showed APA, with 425 (96.8%) expressing mRNA isoforms with shorter 3’UTRs at 48h PHS than in the no heat shock starting condition (Figure 1E). 72hr PHS testes, which have abundant mid-stage apolar spermatocytes, showed APA at 376 genes (279 already detected at the 48hr time point) compared to the no heat shock starting condition. Cumulatively, over the initial 72 hours of the time course, the genome-wide 3’ Seq analysis identified 565 genes that produced mRNA isoforms with a difference in the position of the most abundant 3’ end cleavage site in the 3’UTR in testes enriched for spermatocytes compared to in testes filled with proliferating spermatogonia but lacking spermatocytes (Figure 1F). The time course analysis indicated that most of the APA events occurred at the young spermatocyte stage, with the vast majority (94% - 531/565) resulting in the expression of mRNA isoforms with shorter 3’UTRs as male germ cells differentiate (Figure 1D-G). The 3’UTR shortening due to APA observed in the time course analysis was not a consequence of subjecting flies to heat shock, as similar shortened 3’UTRs were observed in 3’Seq analysis of testes mutant for the spermatocyte specific transcription regulator *aly^-/-^,* which are filled with late-stage spermatocytes (Figure 1B-D, bottom rows). Likewise, analysis of single nuclear RNA-Seq (snRNA-Seq) data from testis produced by the Fly Cell Atlas Consortium (Li et al. 2022) confirmed 3’UTR shortening in spermatocytes compared to spermatogonia for 90% (480 of the 531 genes) detected in our 3’Seq analysis of testes in the differentiation time course (Figure S3).

Taken together, our global 3’Seq analysis of the differentiation time course and the 10X snRNA-Seq results show that a selected set of genes produce mRNA isoforms with shortened 3’UTRs due to utilization of an alternative, more proximal site at which to make the 3’end cut on nascent transcripts after male germ cells undergo differentiation from spermatogonia into spermatocytes. For most genes exhibiting a switch in 3’UTR length due to APA, the short 3’UTR isoform reached its peak level by 48h PHS, corresponding to early spermatocytes at the polar stage (Figure 1G). At 72h PHS, the relative detection of many of the spermatocyte-expressed shorter 3’UTR isoforms decreased, perhaps in part due to the accumulation of new *bam^-/-^* spermatogonia in the *bam^-/-^;hs-Bam* testes by 72hr PHS. For some genes subject to APA, the level of the short 3’UTR isoform continued to increase, possibly due to a substantial increase in levels of transcription from the locus by 72hr PHS. For example, for *orb* (Figure 1D), which encodes an RNA-binding protein with well-known roles in oogenesis (Lantz et al. 1994), the first time point at which a significant number of alternative 3’ cleavage events lead to the production of the short 3’UTR isoform (200nt rather than the 1.2kb 3’UTR present at earlier timepoints) were detected was 32h PHS. The level of expression and the ratio of short rather than long 3’UTR *orb* mRNA isoforms increased at the later time points (48h and 72h PHS). Again, similar short 3’UTR transcripts predominated in testis from flies that had not been subjected to heat shock, where the testes were filled with arrested late spermatocytes due to loss of function of the meiotic arrest gene *aly* (Figure 1D).

### *cis*-regulatory elements differ at proximal versus distal cleavage sites

The set of genes subject to 3’UTR shortening due to APA in early spermatocytes was enriched for weak polyadenylation signal (PAS) motifs (Retelska et al. 2006) near the proximal cut site paired with a strong or canonical PAS motif near the distal cut site. Over half (54.2%) of the 531 genes scored as producing mRNA isoforms with a long 3’UTR in testes enriched for spermatogonia but a short 3’UTR in testes enriched for spermatocytes had either a non-canonical or no known PAS just upstream of the main proximal cleavage site paired with a stronger or canonical PAS just upstream of the main distal cleavage site (Figure 2A-C). *De novo* motif analysis by MEME of sequences extending from the distal cleavage site to 50 bp upstream for the genes that undergo alternative 3’ end cleavage identified the canonical PAS (AAUAAA) (Proudfoot and Brownlee 1976) as the most enriched (e-value 6.2e-21) motif relative to a background data set composed of 3’ cleavage sites of genes that did not undergo APA (Figure 2A). As expected, the position of the PAS motif was around 28 nt upstream of the cleavage site (Proudfoot and Brownlee 1976). Mapping the AAUAAA motif to the genome (AATAAA – sense strand only) revealed that the canonical PAS was present within 50 nt upstream of the distal cleavage site in 65.7% of the genes that produce transcripts with long 3’UTRs in *bam* mutant testes but short 3’UTRs later in the time course (Figure 2B). In contrast, only 22.8% of the genes identified as undergoing stage-specific APA in testes leading to mRNAs with shorter 3’UTRs in spermatocytes had the canonical PAS sequence AATAAA within 50nt upstream of the proximal cleavage site (Figure 2B). In 80% of APA genes, the proximal cut site instead had one of the twelve non-canonical (Retelska et al. 2006), weaker variants of the PAS just upstream (Figure S4A). Analysis of sequences downstream of the proximal and distal cleavage sites in both cases showed enrichment of the expected G/U-rich motif recognized by cleavage factor CstF64 (Figure S4B) (MacDonald et al. 1994).

**Figure 2.**
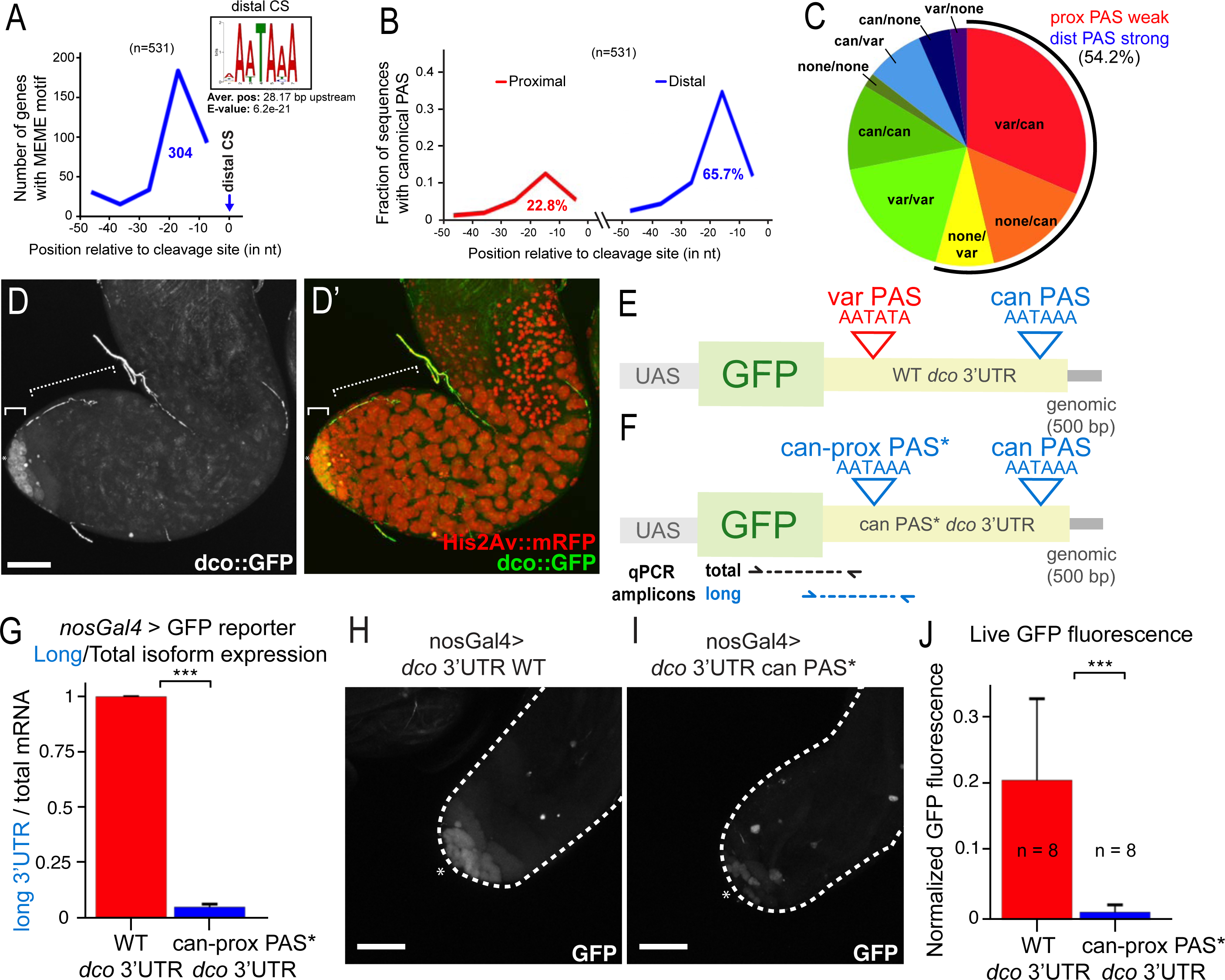
Strength of PAS at proximal site influences stage-specific differential 3’ end formation. (A) Top MEME motif enriched in 50 nt region upstream of distal cleavage sites (CSs) of 531 genes that undergo alternative 3’ cleavage to produce transcripts with shorter 3’ UTRs in later stages of the hsTC, compared to a background of similar 50 nt regions upstream of the 3’ end for genes that do not undergo alternative 3’ cleavage. Plot of the abundance of the MEME enriched sequence along the 50 nt region upstream of the distal CS with number of the APA transcripts that have the MEME motif 10-40 nt upstream of the distal cleavage site in blue. Position of the canonical PAS (AATAAA) upstream of the proximal (red line) or distal (blue line) cleavage sites in genes identified as undergoing the stage specific APA, with percent of the APA transcripts that have a canonical PAS 10-40 nt upstream of the respective cleavage sites. (B) Arrangement of canonical, variant or no PAS motif upstream of the proximal and distal cleavage site (indicated as ‘proximal/distal’) in genes that undergo the stage-specific APA. (Black arc) 54% of the APA genes have a stronger PAS associated with the distal cleavage site than with the proximal cleavage site. (D-D’) Live fluorescent images of testes from *Drosophila* containing a GFP-tagged 3^rd^ copy Fosmid transgene for *dco* (green) and mRFP-tagged His2Av (red). (E, F) Diagram of the paired *dco* 3’ UTR reporter constructs: UAS element (light grey) to drive cell type-specific expression in spermatogonia under the control of *nos-Gal4*; coding region for destabilized GFP (light green); genomic DNA encoding *dco* 3’ UTR (light yellow) plus 500 bases downstream of the distal 3’ cleavage site (dark grey). (E) WT: wildtype *dco* 3’ UTR with the proximal (variant sequence) and distal (canonical sequence) polyadenylation signal (PAS) indicated by a red and blue triangle, respectively. (F) can PAS* *dco* 3’ UTR: same construct as above but with a single nucleotide change in the *dco* 3’ UTR that converts a variant PAS (AATATA) into the canonical PAS (AATAAA) 23 nt upstream of the proximal cleavage site. (G) RT-PCR ratio of reporter mRNA isoform with long 3’ UTR to total reporter mRNA produced in testes from indicated reporters expressed in early germ cells under the control of *nos-Gal4* as measured by qRT-PCR using primer pairs indicated in (F). WT *dco* 3’UTR reporter long/total ratio was set to 1. Error bar indicates s.d. of at least 3 independent biological replicates. (H and I) Native GFP fluorescence for the respective reporters expressed at the apical tip of testes (indicated by an asterisk) under control of *nos-Gal4*. (J) Quantification of GFP fluorescence in Z-stacks through apical tips of live testis, normalized to His2Av::mRFP expression. Asterisk: hub; solid bracket: spermatogonia; dashed bracket: spermatocytes. Scale bars: 50 microns. Statistical significance determined by two-tailed student’s t-test (*** indicates *p*-value < 0.001).

To test whether the pairing of weak proximal PAS with strong distal PAS was important for the stage-specific APA that results in the expression of mRNA isoforms with long 3’UTRs in spermatogonia but short 3’UTRs in spermatocytes, we constructed reporter transgenes containing the 3’UTR region from *dco,* which shows APA as male germ cells differentiate from spermatogonia to spermatocytes (Figure 1C). Analysis of expression of a GFP-tagged *dco* fusion protein encoded by a large Fosmid-based transgene containing over 23kb of genomic DNA, including 5kb upstream and 10kb downstream of *dco* (FlyFos TransgeneOme (fTRG) project (Sarov et al. 2016)) showed that the Discs overgrown (Dco) protein was strongly expressed in the cytoplasm of germline stem cells (GSCs), gonialblasts (Gbs) and spermatogonia at the testis apical tip (Figure 2D - solid bracket). However, expression of Dco-GFP from the Fosmid transgene was abruptly downregulated when germ cells became spermatocytes (Figure 2D - dotted bracket). The 3’UTR region of the *dco* locus has a PAS variant (AATATA) upstream of the proximal cleavage site and a canonical PAS (AATAAA) upstream of the distal cleavage site. We constructed a matched pair of reporter transgenes for the *dco* locus that each contained the inducible UASt promoter (5X UAS repeats) upstream of the coding region for a destabilized GFP (Li et al. 1998) followed by genomic DNA encoding the entire long 3’UTR sequence plus 500 bases of genomic DNA from the *dco* locus downstream of the distal cleavage site (Figure 2E, F). In the wild-type reporter, the 3’UTR contained the variant PAS AATATA 22 nt upstream of the proximal cleavage site and the canonical PAS AATAAA 33 nt upstream of the distal cleavage site, as in the endogenous *dco* locus (Figure 2E). The mutated transgenic reporter *can-prox PAS\** was identical except for one base change that altered the variant proximal PAS AATATA to the canonical PAS AATAAA (Figure 2F). The two transgenes were stably inserted into the same genomic attP landing site in separate *Drosophila* lines, and transcription of the reporters was induced specifically in germline stem cells and early spermatogonia using the *nanos-Gal4* expression driver to assess the effect of mutating the proximal PAS site on mRNA 3’ end processing in spermatogonia.

Analysis of expression of mRNA isoforms with long vs. short 3’UTRs *in vivo* from the two *dco* reporter transgenes transcribed in GSCs and early spermatogonia under control of *nos-Gal4* revealed that the single nucleotide change converting the variant PAS upstream of the proximal *dco* cleavage site to the canonical PAS was sufficient to allow 3’UTR shortening in spermatogonia. Analysis by qRT-PCR to detect the ratio of mRNA from the GFP reporter transgene processed with the predicted long *dco* 3’UTR relative to total GFP reporter mRNA revealed that for the reporter mutated to have the canonical PAS upstream of the proximal cleavage site (*can-prox PAS\**), the ratio of the long 3’UTR to total mRNA was 22-fold decreased relative to the wild-type reporter (Figure 2G). The total level of mRNA expressed from the *can-prox PAS\** reporter did not decrease relative to the wild-type reporter (Figure S4D). Together, these data suggest that, for the *dco* locus, an exact match to the canonical PAS signal AATAAA is required in spermatogonia for making the 3’ end cut that terminates nascent transcripts.

Imaging of GFP fluorescence in testes from flies carrying the reporters expressed under the control of *nanos-Gal4* showed much lower levels of GFP in spermatogonia from the mutated transgene (Figure 2H), which produced transcripts with the short 3’UTR (Figure 2G), even though the two reporters had the same protein-coding region sequence (Figure 2E, F). Although the level of GFP fluorescence varied from cyst to cyst, as has been observed by others when the expression is driven under the control of Gal4/UAS (Skora and Spradling 2010), the reporter with the wild-type *dco* 3’UTR showed substantial expression of GFP in proliferating germ cells at the tip of the testis (Figure 2H) while the reporter mutated to contain a canonical PAS upstream of the proximal cleavage site (can-prox PAS*) resulted in a 20-fold decrease in GFP fluorescence, with weak and variable expression detected in only a few germ cells at the testis tip (Figure 2I, J). Together, these data suggest that sequences in the long *dco* 3’UTR expressed in spermatogonia may facilitate translation of the *dco* mRNA into protein and that removal of these sequences in young spermatocytes by cleavage of nascent transcripts at the more proximal cut site may lead to an abrupt shut down of Dco protein expression.

### 3’UTR shortening by APA correlates with changes in protein expression

The changes in Dco protein expression as spermatogonia become spermatocytes led us to analyze the dynamics of protein expression in other cases of genes subject to 3’UTR shortening due to APA where reagents to follow protein expression were available. Expression of the LolaF protein, encoded by mRNA isoforms resulting from 3’UTR shortening due to stage-specific APA starting in young spermatocytes (Figure 3A), showed regulatory dynamics reciprocal to the Dco protein. Immunostaining of wild-type testes with antibodies specific for the LolaF protein (Zhang et al. 2003) detected no signal in spermatogonia at the testis apical tip but abundant nuclear LolaF protein in spermatocytes (Figure 3B). The timing of 3’UTR shortening by APA correlated well with the onset of protein expression. Analysis by 3’RACE of *lolaF* transcripts from testes at different stages of the heat shock time course confirmed that the change in the 3’end cut site for *lolaF* occurred soon after the onset of spermatocyte differentiation, with the short 3’UTR isoform appearing by 32hr PHS and persisting as the predominant 3’RACE product through 48hr PHS (Figure 3C). Strikingly, although the *lolaF* mRNA isoform with a long 3’UTR was abundantly expressed in *bam* mutant testes based on our 3’Seq and 3’RACE data (Figure 3A, C), LolaF protein was not detected by immunostaining in *bam^-/-^;hs-Bam* mutant testes either before heat shock or 16hr PHS (Figure 3D, E). However, by 32 hours PHS, correlating with the appearance of the short 3’UTR mRNA isoform, LolaF protein detected by immunofluorescence staining was sharply upregulated in the differentiating germ cells (Figure 3F, G).

**Figure 3.**
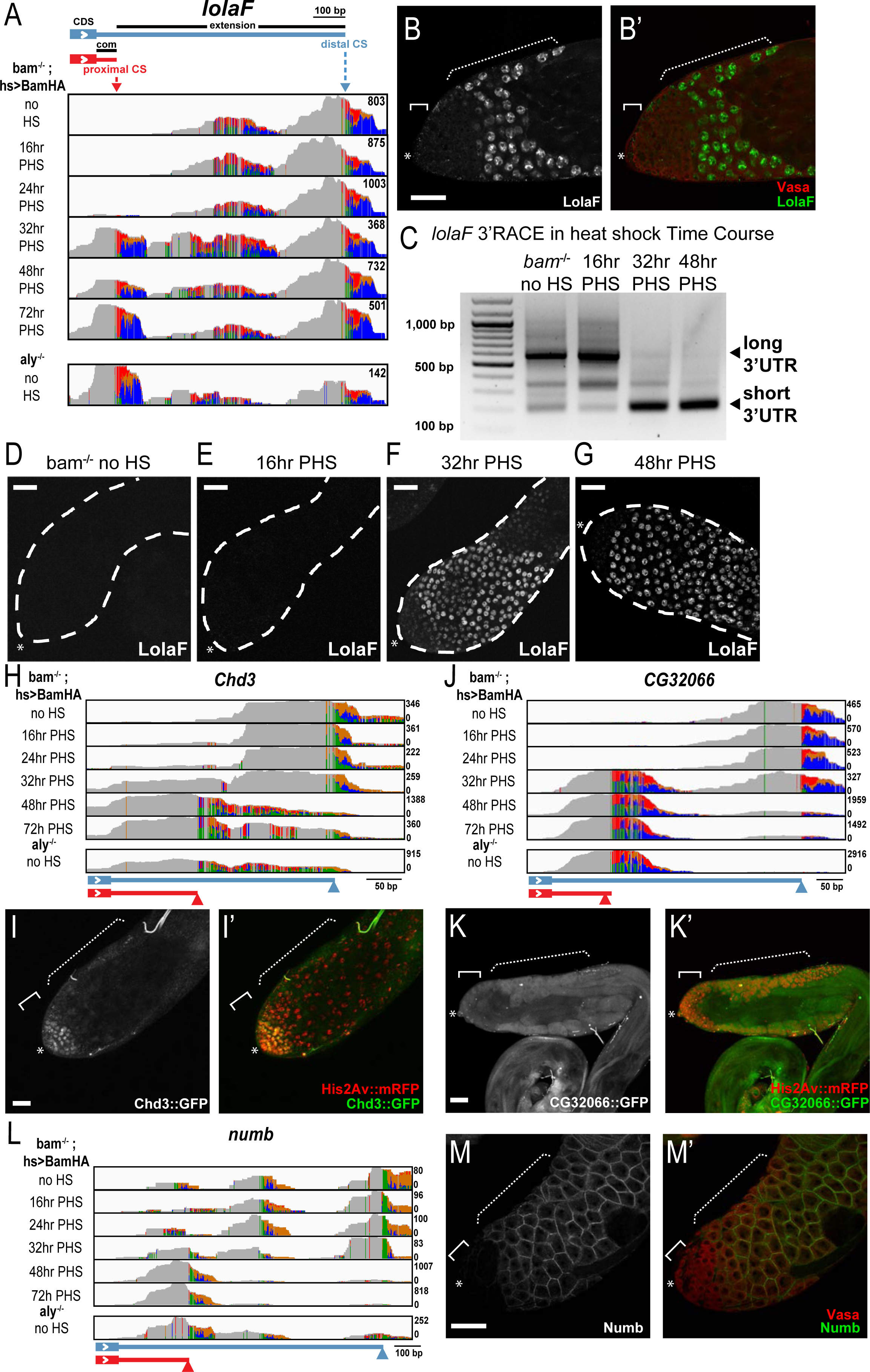
Changes in protein expression accompany the shortening of 3’UTRs. (A) 3’ Seq tracks of testis mRNA from one of two biological replicates plotted on the 3’ genomic region of *lolaF* from the *bam* hsTC at indicated times PHS, as well as from and *aly* mutant testes. (B, B’) Immunofluorescence image of the apical region of a wild-type testis stained with anti-LolaF (white/green) and anti-Vasa (red) antibodies. (C) 3’RACE of *lolaF* from *bam* hsTC fly testes at indicated time points showing the expression of long and short 3’UTR isoform. (D-G) Immunofluorescence images containing the apical third of testes from *bam*^-/-^*; hs-Bam* flies with no heat shock or 16, 32, or 48hrs PHS stained with anti-LolaF antibody. (H, J, L) 3’ Seq tracks from one of two biological replicates from testes from *bam* hsTC flies at indicated times PHS, as well as from *aly* mutant testes, plotted on the 3’ genomic regions for (H) *Chd3*, (J) *CG32066*, (L) *numb.* Bottom: Proximal (red arrowhead) and distal (blue arrowhead) cleavage sites were predicted from the most highly used cleavage site in *bam^-/-^; hs-Bam* testes without heat shock and 48hr PHS, respectively. (I, K, M) Native fluorescence from GFP (white/green) and mRFP (red) in live mount testes from flies carrying His2Av-mRFP and 3^rd^ copy Fosmid-based GFP tagged reporters for (I, I’) Chd3-GFP, (K, K’) CG32066-GFP. (M, M’) Immunofluorescence staining of testis tip with anti-Numb (white/green) and anti-Vasa (red). Asterisk: hub; solid bracket: spermatogonia; dashed bracket: spermatocytes. Scale bars: 50 microns.

Analysis of expression of GFP-tagged fusion proteins encoded by large, Fosmid based genomic transgenes available through the FlyFos TransgeneOme (fTRG) project showed several other cases of genes that encode mRNA isoforms with long 3’UTRs in spermatogonia but short 3’UTRs in spermatocytes where expression of the protein changed dynamically with male germ cell differentiation. Like Dco-GFP, the GFP tagged fusion protein encoded by a Fosmid transgene for Chd3 was expressed in spermatogonia but not detected in young spermatocytes, correlating with 3’UTR shortening by stage-specific APA (Figure 3H, I). In contrast, the GFP-tagged fusion protein encoded by a Fosmid transgene for the protein encoded by CG32006 (Figure 3J, K) was, like LolaF, strongly upregulated in spermatocytes compared to spermatogonia). In addition, immunofluorescence staining with antibodies against Numb (O’Connor-Giles and Skeath 2003) showed a lacework pattern of Numb protein localized at the periphery of spermatocytes but did not detect the protein in spermatogonia (Figure 3L, M). The reciprocal dynamics of protein expression for the genes assessed suggest that a single molecular event, developmentally regulated APA, may trigger different changes in expression of the encoded proteins, with some going from OFF in spermatogonia to ON in spermatocytes and others from ON in spermatogonia to OFF in spermatocytes, presumably depending on the *cis*-regulatory sequences present in the different mRNA isoforms.

### Polysome fractionation suggests 3’UTR shortening correlates with switches in translation activity

To investigate globally whether changes in translation state may be a widespread consequence of stage-specific 3’UTR shortening by developmentally regulated APA, we carried out polysome fractionation followed by 3’Seq. Lysates of testes from *bam^-/-^;hs-Bam* flies 24hr (enriched in transcripts with long 3’UTRs), 48hr (enriched in transcripts with short 3’UTRs), or 72hr PHS were cleared of nuclei, layered on sucrose gradients, centrifuged to separate fractions with zero, few, or many ribosomes, then assessed by 3’Seq to score co-migration of mRNA isoforms (Figures 4A, B). The results revealed that shortening of the 3’UTR by stage-specific cleavage of transcripts at the proximal site frequently correlated with dramatic differences in the position at which the two mRNA isoforms migrated in the polysome profiles. For 508 of the 531 genes we identified as undergoing APA to produce mRNA isoforms with shorter 3’UTRs as male germ cells differentiate from spermatogonia to spermatocytes, the depth of 3’Seq from the sucrose gradient fractions containing free RNA, 40S or 60S ribosomal subunits, 80S ribosomes, 2-3 ribosomes, or 4 or more ribosomes was sufficient to obtain information for how the long 3’UTR mRNA isoform behaved in extracts from 24h PHS testes and how the short 3’UTR mRNA isoform behaved in extracts from the 48h PHS testes samples. For one quarter (124/508) of the genes identified as undergoing stage-specific APA leading to 3’UTR shortening with male germ cell differentiation, polysome fractionation suggested that the long 3’UTR isoform expressed at early time points was translationally silent: the long 3’UTR isoform was detected in the ribosome-free, 40S, and/or 60S ribosomal subunit-containing fractions but did not substantially co-migrate with the fractions containing one, 2-3, or 4 and more ribosomes in testis extracts from 24h PHS (Figure 4C). For 50 of these “Long 3’UTR OFF” genes, the short 3’UTR isoform expressed from the same gene at 48h PHS was detected in the fractions containing 80S monosomes, 2-3 ribosomes, or 4 and more ribosomes, indicating a substantial difference in behavior of the mRNA isoforms (Figure 4D).

**Figure 4.**
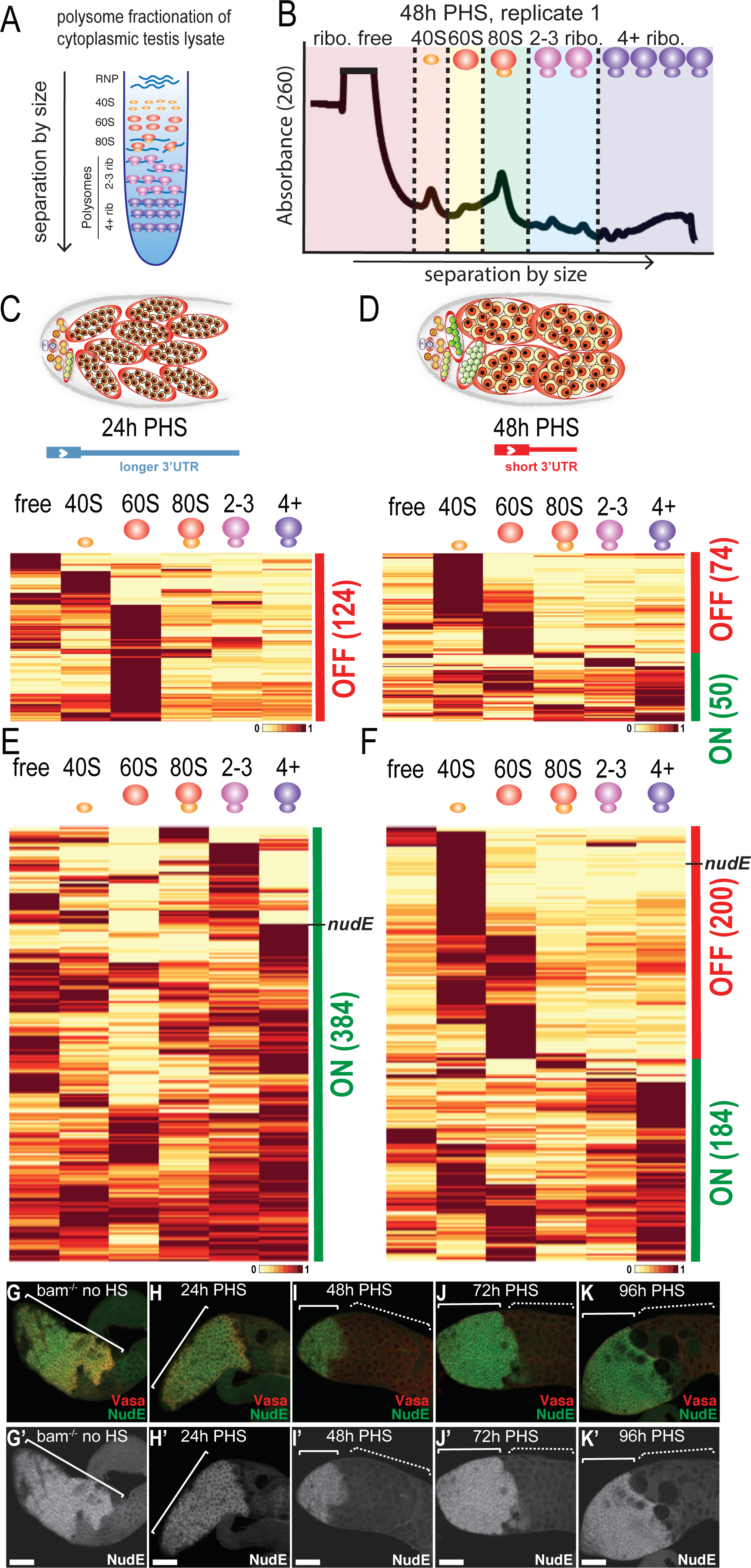
3’ Seq following polysome fractionation reveals widespread differences in migration of transcripts with long 3’UTRs at 24hr PHS vs their short 3’UTR isoforms at 48hr PHS. (A) Diagram of polysome fractionation by centrifugation: transcripts occupied by more ribosomes sediment further down in the polysome profile. (B) Absorbance at 260 from one of two replicates of the polysome profile of testis extracts from 48h PHS, indicating separation of the 40S, 80S, and multiple polysome peaks. ROYGBV colors (with dashed lines as boundaries) indicate the polysome fractions that were combined before 3’ Seq. (C and E) 3’ Seq from polysome profiling from 24hr PHS testes, plotting relative level of the long 3’UTR isoform detected across the different polysome fractions (in each column) for genes that undergo the stage-specific 3’UTR shortening due to APA (in each row). (C) Heatmap of the 124 long 3’UTR mRNA isoforms that predominantly comigrated with polysome fractions lighter than the 80S at the 24hr PHS timepoint. (E) Heatmap of the 384 long 3’UTR mRNA isoforms that comigrated with the 80S and/or polysomes at the 24hr PHS timepoint. (D and F) 3’ Seq from polysome profiling of 48hr PHS testes, plotting relative level of the short 3’UTR isoform detected across the different polysome profile fractions for genes that undergo the stage-specific 3’UTR. (D) Heatmap of distribution based on 3’ Seq of the polysome fractions from 48hr PHS testes of the short 3’UTR mRNA isoforms from the 124 genes grouped in (C). (F) Heatmap of distribution based on 3’ Seq of the polysome fractions from 48hr PHS testes of the short 3’UTR mRNA isoforms from the 384 genes from (E). Heatmap: white/yellow indicates low expression of indicated transcript; dark red indicates higher expression of the transcript. *nudE* transcript indicated in black. (G-K) Immunofluorescence images of testes from *bam ^Δ86^*^/1^; hs-Bam flies with no heat shock or 24, 48, 72 or 96 hrs PHS stained with anti-nudE antibody (white/green) and anti-Vasa (red). Solid bracket: spermatogonia; dashed bracket: spermatocytes. Scale bars: 50 microns.

For three quarters (384/508) of the genes identified as undergoing stage-specific APA leading to 3’UTR shortening with male germ cell differentiation, the isoform with the long 3’UTR expressed at 24h PHS was detected in the fractions with one (80S), 2-3, or 4 and more ribosomes (Figure 4E). Strikingly, for over half (200/384), the isoform with the short 3’UTR expressed from the same gene was almost exclusively present in the free, 40S and/or 60S fractions in testis extracts from 48h PHS, indicating translational repression (Figure 4F). Immunofluorescence staining of testes with available antibodies against the protein product of one of these genes, NudE (Wainman et al. 2009), revealed that the protein is strongly expressed in spermatogonia but is abruptly downregulated in the young spermatocytes differentiating in testes 48hr PHS (Figure 4G-I), as predicted from the migration behavior of the mRNA isoforms upon polysome fractionation (Figure 4E, F, S6A). Levels of immunofluorescence signal for NudE protein remained low in mid-stage spermatocytes present at 72hr PHS and the maturing spermatocytes present at 96hr PHS (Figure 4J, K: dotted brackets) although high levels of immunofluorescence were detected in the *bam^-/-^* spermatogonia accumulating at the apical tip of the testis (Figure 4J, K: solid brackets).

Together, the results of polysome fractionation followed by 3’Seq suggest that the stage-specific switch in 3’UTR processing that forms the 3’ end of nascent transcripts can result in dramatic changes in the interaction of the resulting mRNA isoforms with ribosomes. If this reflects changes in translation state, then developmentally regulated APA may allow sharp shifts in expression of many proteins as cells move from a proliferation program to onset of differentiation, some from ON → OFF and others from OFF → ON.

### Reactivation of translation in later germ cell stages

Some mRNA isoforms with a short 3’UTR due to APA that were translationally repressed at 48h PHS began to co-migrate with monosomes by 72hr PHS. For 199 of the 200 genes where the long 3’UTR mRNA isoform migrated in fractions containing monosomes or higher numbers of ribosomes in 24hr PSH testis extracts while the short 3’UTR isoform migrated in sub-monosomal fractions at 48hr PHS (Figure 4C, D), there was sufficient read depth in fractions from polysome gradients of 72hr PHS testis samples to follow the migration behavior of the short 3’UTR isoforms at the later time point. For 69% of these 199 genes, the short 3’UTR mRNA isoform showed significant enrichment in fractions co-migrating with the 80S monosome (106/199; 53%) and/or polysomes (33/199; 16%) in lysates of testes taken 72hr PHS (Figure 5A, B). For the remaining 31%, the short 3’UTR mRNA isoform remained associated with sub-monosomal fractions at 72hr PHS, suggesting these transcripts remain translationally repressed.

**Figure 5.**
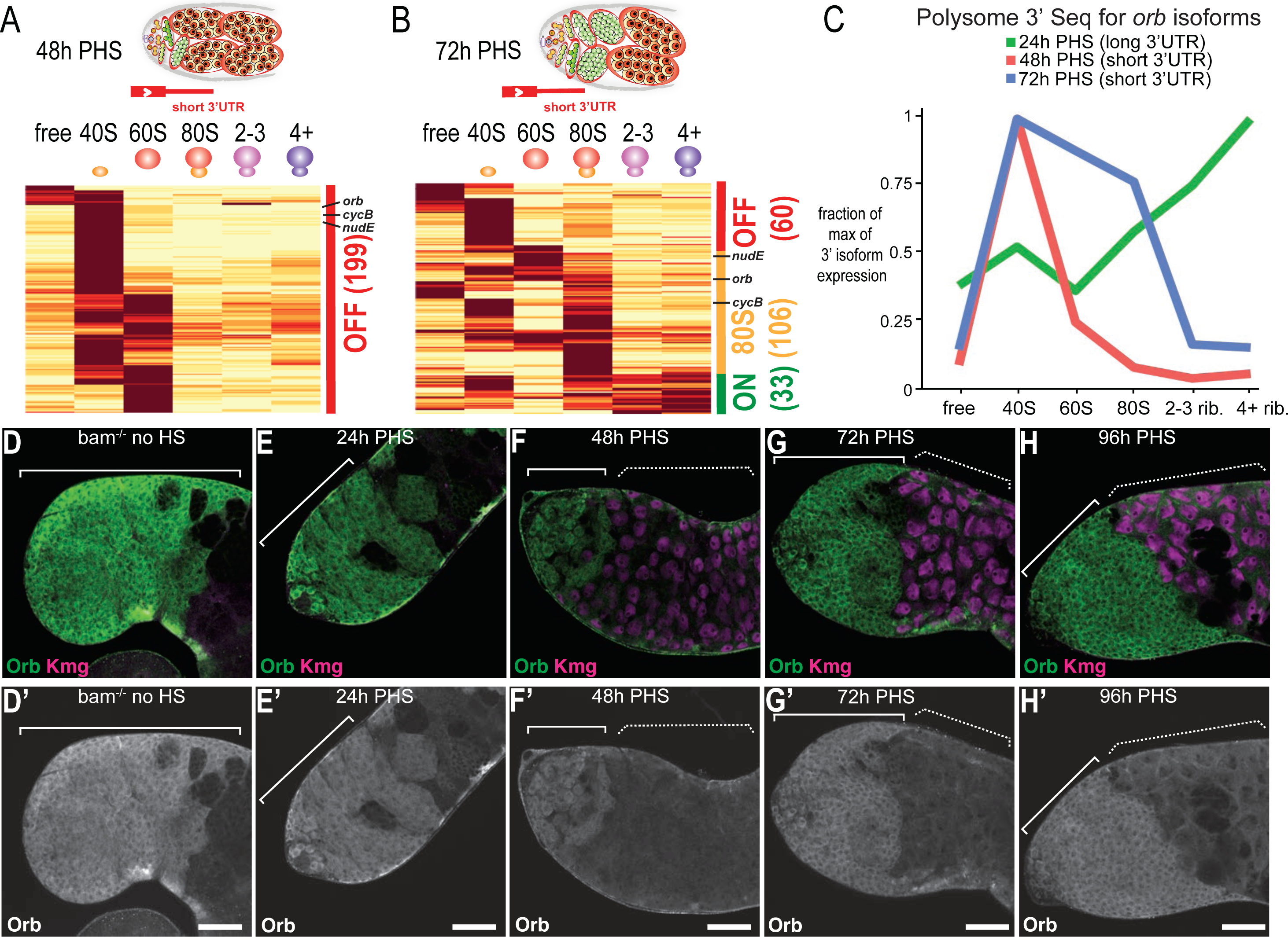
Dynamic changes in polysome profile of short 3’UTR isoforms as spermatocytes mature revealed by 3’ Seq polysome profiling of 72h PHS testes. (A) Heatmap of distribution in the polysome gradient based on 3’ Seq of 199 of the short 3’UTR mRNA isoforms that predominantly comigrated with fractions lighter than the 80S monosome at the 48hr PHS timepoint from Figure 4F. (B) Heatmap of distribution in the polysome fractions from 72hr PHS testes of the same 199 short 3’UTR mRNA isoforms shown in A. (C) Line graph of relative levels of the long 3’UTR *orb* isoform (green, 24h PHS) and short 3’UTR *orb* isoform (red, 48h PHS; blue, 72h PHS) in the indicated polysome fractions. (D -H) Immunofluorescence images of apical regions of testes from *bam ^Δ86^*^/1^; hs-Bam flies with no heat shock or 24, 48, 72 or 96 hrs PHS stained with anti-Orb antibody (white/green) and antibody against the spermatocyte marker Kmg (magenta). Solid bracket: spermatogonia; dashed bracket: spermatocytes. Scale bars: 50 microns.

*cyclinB1* is one already known example of a gene that encodes an mRNA isoform with a long 3’UTR that is translated in spermatogonia and a short 3’UTR mRNA isoform that is translationally repressed in young spermatocytes but reactivated to produce protein in mature spermatocytes before the G2/M transition of meiosis I (White-Cooper et al. 1998; Baker et al. 2015). Consistent with this, the short 3’UTR *cyclinB1* mRNA isoform expressed in spermatocytes migrated mostly in the 40S fraction in extracts from testis 48hr PHS but was substantially present in the 80S monosomal fraction by 72hr PHS (Figure 5A, B, S6B).

Antibody staining over the time course of spermatocyte maturation confirmed an additional example where polysome fractionation suggested that the short 3’UTR mRNA isoforms were not translated in young spermatocytes but began to produce protein as spermatocytes mature. For *orb*, the long 3’UTR mRNA isoform expressed in testes filled with proliferating spermatogonia (Figure 1D, G) comigrated with polysomes in sucrose gradients of extracts of 24h PHS testes (Figure 5C, green line). In contrast, the *orb* short 3’UTR mRNA isoform expressed in testes filled with differentiating spermatocytes (Figure 1D, G) was detected predominantly in the 40S fraction in sucrose gradients at 48h PHS testes (Figure 5C, red line). However, in extracts from 72h PHS testes the *orb* short 3’UTR mRNA isoform showed a substantial increase in signal in the 80S monosome fraction (Figure 5C, blue line). Consistent with the behavior of *orb* mRNA isoforms in the polysome gradient fractionation assay, immunofluorescence staining with anti-Orb antibodies revealed Orb protein expressed in proliferating progenitor cells filling testes from *bam^-/-^;hs-Bam* flies without heat shock or at 24h PHS (Figure 5D, E), but decreasing sharply in the early spermatocytes (marked by expression of Kumgang (Kmg) (Kim et al. 2017)) differentiating in testes 48h PHS (Figure 5F), consistent with the dramatic shift from co-migration with polysomes at 24h PHS to predominantly with the 40S fraction by 48h PHS (Figure 5C). However, consistent with the detection of some *orb* short 3’UTR mRNA isoform in the 80S monosomal fraction by 72hr PHS (Figure 5A-C) indicating possible reactivation of translation, testes from 96hr PHS flies showed an increase in immunofluorescence signal for Orb protein in late-stage spermatocytes about to initiate the meiotic divisions (Figure 5H, dotted bracket).

## Discussion

Here we show that a stage-specific change in 3’end cleavage site choice leading to the production of mRNA isoforms with short rather than long 3’UTRs can trigger dramatic and widespread changes in the suite of proteins expressed as cells progress from one developmental stage to the next in a differentiation lineage. In some cases, 3’UTR shortening by developmentally regulated APA correlates with a switch from protein expression in proliferating spermatogonia to repression of translation in young, differentiating spermatocytes (ON -> OFF), while for other genes the switch is from no protein expressed in spermatogonia to protein accumulation in spermatocytes (OFF -> ON). Thus, a single molecular event, developmentally regulated APA, can trigger changes in the expression of many proteins, likely depending on the *cis*-regulatory sequences present in the long 3’UTR isoform expressed by individual genes.

Our findings in the *Drosophila* male germline adult stem cell lineage may have implications for other cases where widespread APA has been documented. For example, the production of mRNA isoforms with long 3’UTR extensions noted in the nervous system in a variety of animals (Hilgers et al. 2011; Miura et al. 2013; Vallejos Baier et al. 2017) may allow translational repression of specific mRNAs unless they are located at activated synapses (An *et al*. 2008; Andreassi *et al*. 2010; Yudin *et al*. 2008). Conversely, the widespread expression of mRNA isoforms with short 3’UTRs due to APA in proliferating cancer cells (Sandberg et al. 2008; Mayr and Bartel 2009; Masamha et al. 2014; Xia et al. 2014) may contribute to cancer progression by allowing abnormal expression of oncogenic proteins usually translationally repressed via sequences in the longer 3’UTRs expressed in normal cells. It is possible that the mechanisms that normally ensure proper 3’cut site selection in many mammalian cell types are anti-oncogenic, much as is the proper function of genes involved in DNA repair such as BRCA1 and BRCA2, with defects that result in abnormal use of more proximal, less favored PAS signals selected for in tumor cells because they allow promiscuous expression of oncogenic proteins. We note that although 3’ UTR shortening in cancer cells has been correlated with proliferation, the trend is opposite in the adult stem cell lineage we document: the genes subject to stage-specific APA in the *Drosophila* male germline mainly produce mRNA isoforms with long 3’UTRs in proliferating spermatogonia but shortened 3’UTRs once differentiation into spermatocytes begins.

Our results in the adult stem cell lineage differentiation agree with the findings in HEK 293T and five other human cell lines showing that alternative processing of 3’UTRs broadly influences translation (Floor and Doudna 2016; Fu et al. 2018). Changes in 3’UTR length may be associated with miRNA-mediated mRNAs regulation as seen in cancer, where the shortening of transcripts allows escape from miRNA targeting (Mayr and Bartel 2009). Also, 3’UTRs may be bound at specific sequences by numerous RBPs that can influence the stability, localization, and translation of mRNAs, as is the case of AU-rich binding proteins known to repress translation (García-Mauriño et al. 2017).

Why might normal cells use such an APA mechanism to regulate changes in protein expression as they advance from one state to the next in a developmental progression? As discussed in the introduction, the switch from precursor cell proliferation to the onset of terminal differentiation involves dramatic changes in cellular programs that must be cleanly executed. Developmentally regulated APA may provide a mechanism to rapidly turn off the expression of specific proteins for the prior proliferation program and initiate the expression of proteins involved in the onset of differentiation to facilitate clean and sharp transitions between developmental states. Since APA occurs on nascent transcripts, 3’UTR shortening that removes sequences that instruct translational repression of the mRNA may facilitate the rapid onset of expression of the encoded protein without having to await chromatin opening, formation of a preinitiation complex, then initiation and elongation of transcripts as would be required in turning on a new transcription program.

Results of network and gene enrichment analyses were consistent with the idea that an APA-based mechanism to shut off protein expression may aid transition from mitotic proliferation to onset of differentiation. Functional network analysis and Gene Ontology (GO) enrichment analysis using STRING of the 531 genes that undergo APA resulting in shorter 3’ UTRs in differentiating spermatocytes revealed enrichment in the categories of Biological Process GO term “Cell cycle” and Cellular Component GO term “Polycomb Group (PcG) protein complex” (Figure S7A, highlighted in red and blue, respectively). Similar analysis of the 200 genes that undergo 3’UTR shortening where polysome profiling suggested a switch from active translation of the long 3’UTR isoform in proliferating spermatogonia (24h PHS) to a translationally inactive state of the short 3’UTR isoform in differentiating spermatocytes (48h PHS) showed enrichment of the Biological Process GO term “Cell cycle process” (Figure S7B, highlighted in red). Enrichment of cell cycle regulators in the set of APA genes predicted to switch from protein expression ON in spermatogonia to OFF in spermatocytes is consistent with previous studies showing that other cell cycle regulatory proteins, PCNA (Insco et al. 2009) and DREF (Angulo et al. 2019), are abruptly downregulated after completion of pre-meiotic S Phase and entry into meiotic prophase in the *Drosophila* male germline. PcG components were also enriched among the 531 genes that undergo 3’UTR shortening by APA, although polysome fractionation results suggested that some may switch from ON -> OFF, whole other may switch from OFF -> ON. In addition to the APA genes noted here, protein expression of the PRC2 components E(z) and Su(z)12 (Chen et al. 2011) have been shown to switch from ON -> OFF as spermatogonia differentiate into spermatocytes. Together the results suggest that there may be widespread changes in protein expression for PcG components during germline differentiation, perhaps reflecting a duty shift between homologs.

An APA-based mechanism to shut off protein expression as precursor cells cease mitosis and initiate differentiation may be especially important during spermatogenesis, as many cell cycle regulators and that could potentially be deleterious early in meiotic prophase are called into action later for spermatocytes to enter and execute the meiotic divisions. In addition, many genes involved in later stages of male germ cell differentiation must be expressed in spermatocytes to endow spermatids with mRNAs they will translate later. Our 3’Seq analysis of polysome gradient fractions from testis filled with young (48hr PHS) or maturing (72hr PHS) spermatocytes revealed that many mRNA isoforms with shortened 3’UTRs may become translationally derepressed as spermatocytes mature, moving to migrate with the 80S monosome or polysomal fractions. This dynamic translational regulation following alternative 3’ cleavage may allow differentiating germ cells to repress the expression of specific proteins for a defined but critical period during differentiation onset, while still maintaining transcripts for re-expression of the protein at later stages (Figure 6).

**Figure 6.**
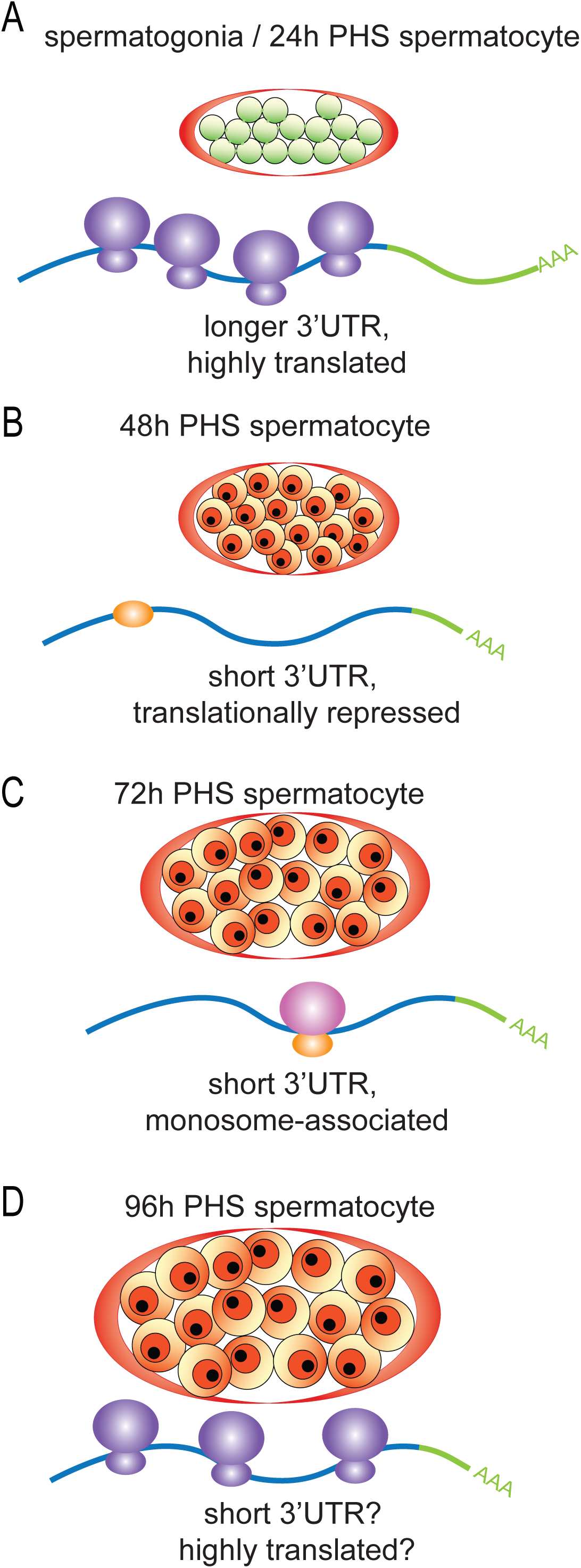
Model of dynamic translational regulation of transcripts from genes subject to APA as *Drosophila* germline progenitors differentiate. Cartoon depictions of germline cysts from (A) 24 hr, (B) 48hr, (C) 72hr, and (D) 96hr PHS testes. Predicted transcript association with ribosomes based on 3’ Seq polysome profiling with the coding region (blue) and 3’UTR (green).

We identified 200 genes subject to stage-specific APA where the long 3’UTR isoform co-migrated with polysomes at 24h PHS but the short 3’UTR isoform expressed at 48hr PHS co-migrated with sub-monosomal fractions, suggesting translational repression. For many of these, the short 3’UTR isoform was enriched in the gradient fractions containing the 40S small ribosomal subunit at 48hr PHS. Co-migration of the short 3’UTR isoform with the 40S or 60S sub-ribosomal fractions may reflect transcripts accumulating at an intermediate step in translational activation (Sokabe and Fraser 2019). Alternatively, the co-migration of many short 3’UTR mRNA isoforms with the 40S small ribosomal subunit at 48h PHS we observed may involve a mechanism where translationally repressed mRNAs are complexed with the 40S ribosomal small subunit, as has been shown in stress granules, regulated by phosphorylation of eIF2 (Panas et al. 2016).

Our combined polysome fractionation and 3’Seq indicated that for half of the ∼500 genes we identified as subject to 3’UTR shortening by stage-specific APA, the mRNA isoforms with short 3’UTRs expressed in young spermatocytes migrated differently in polysome gradient fractions than the mRNA isoforms with long 3’UTRs expressed in spermatogonia, suggesting changes in translation state. However, changes in protein expression upon APA may be even more widespread. Analysis of protein expression for several genes, including *dco, LolaF, Chd3*, *numb,* and *CG32066* (Figure S6C-G), showed dramatic up or down-regulation of protein expression correlating with the timing of 3’UTR shortening by APA even though the long and short 3’UTR mRNA isoforms did not show dramatic changes in migration pattern in polysome gradients. It is possible that in some cases translational repression does not result in changes in the polysome profile of the mRNA, perhaps due to cotranslational degradation of mRNAs targeted by microRNAs, as has been shown for 3’UTR dependent repression of translation of *hid* by the bantam miRNA and *reaper* by miR-2 in *Drosophila* S2 cells (Tat et al. 2016). Additional studies have shown that regulation of *lin-41* by the *let-7* microRNA or of *lin-14* by the *lin-4* microRNA in *C. elegans* (Olsen and Ambros 1999; Stadler et al. 2012) also occurs without changing the polysome profile of miRNA-targeted mRNAs.

Together, our results demonstrate that 3’UTR shortening due to developmentally regulated APA can lead to widespread changes in the suite of proteins expressed at specific steps in a differentiation sequence. Strikingly, protein expression can be either turned from ON to OFF or from OFF to ON, in a gene selective manner, presumably controlled by sequences in the longer 3’UTR expressed in proliferating precursor cells.

## Materials and Methods

### Fly strains and husbandry

*Drosophila* strains were raised on standard molasses medium at 25°C unless otherwise noted. The following *Drosophila* strains were used: *w^1118^* as wild-type; *bam^1^* (McKearin and Spradling 1990); *bam^Δ86^* (McKearin and Ohlstein 1995); *bgcn^1^*, *bgcn^63-44^* (Ohlstein et al. 2000); *bam-Gal4* (Chen and McKearin 2003); *aly^5p^*, *aly^2^* (White-Cooper et al. 2000); *sa^1^* and *sa^2^*(Lin et al. 1996); Tagged FlyFos TransgeneOme (fTRG) (Sarov et al. 2016)stocks were obtained from VDRC.

For the heat shock time course (hsTC), *hs-BamHA/CyO*; *bam^Δ86^,e/TM3,e,Sb* males and *bam^1^,e/TM6b,e,Hu* females were crossed on molasses medium at 22°C, adults removed after 3 days, and after approximately 9 days after crossing late-stage pupae were shifted for 30 min to a 37°C water bath for heat shock, and then incubated at 25°C for 16, 24, 32, 48, 72 or 96 hours before collection of *hs-BamHA/+*;*bam^1^/bam^Δ86^*flies for dissection.

### 3’ Seq Data Analysis

Reads obtained from sequencing on the Illumina Nextseq 500 platform from libraries created with the QuantSeq 3′ mRNA-Seq Library Prep Kit FWD for Illumina (Lexogen) (including oligo dT priming and insert size optimization for shorter read lengths of 50 - 100 nts) were aligned against the *Drosophila* (dm6) genome using BOWTIE version 0.12.8 (Langmead et al. 2009) with a value of e = 5000 to allow for expected mismatches in the poly A tail not encoded in the genome (Figure S1A). Cleavage sites were defined as the location of the first A in a string of at least 15 contiguous As requiring that at least 3 do not match the genome. All cleavage sites reported supported by at least 20 reads. Minor CSs that were within 50 bp of another cleavage site with a higher read count were discarded. Cleavage sites that did not map to annotated Flybase 3’ UTRs (r6.36) or within 500 bp downstream of annotated 3’ UTRs were not analyzed in this study. To prevent the association of cleavage sites in 5’UTRs of neighboring genes with the upstream gene caused by the 500 bp 3’ UTR extension, we did not extend 3’UTRs into a neighboring gene’s 5’ UTR as defined by Flybase. Cleavage sites were assigned to the closest upstream CDS end as defined by Flybase.

To identify genes encoding transcript isoforms arising from differential 3’ end cleavage in the hsTC, we identified the main cleavage site for each transcript in each library, defined as the cleavage site supported by the highest number of reads for each transcript. To call differential 3’ end cleavage of transcripts expressed at different timepoints, we required the peaks to pass criteria on both biological replicates of a given timepoint, and the distance between the main cleavage sites at different timepoints to be greater than 100bp (Figure S1A). Furthermore, we required genes that gave rise to transcripts with different main cleavage sites to have greater than a 4-fold difference in expression level between the two isoforms in *bam*^-/-^; *hs-Bam-HA* flies without heat shock vs 16h, 24h, 32h, 48h or 72h PHS libraries.

To validate the reproducibility of the 3’ Seq protocol in the biological replicates, we plotted the expression of all 3’ cleavage sites expressed in both biological replicates from various timepoints in the hsTC and *aly* mutant testes (Figure S1C-I). The majority of the datasets have high values for the coefficient of determination *R*^2^ as well as the Pearson correlation coefficient *r*, indicating that the 3’ Seq protocol performed on replicate biological tissue from the hsTC testes yielded reproducible results.

### MEME motif analysis

Unbiased motif searches on sequences spanning 50 bases upstream of proximal or distal cleavage sites for genes that undergo APA were performed using MEME (Multiple Em for Motif Elucidation) version 5.4.1 (Bailey and Elkan 1994; Bailey et al. 2015), a discriminative motif discovery algorithm. Control sequences were defined as 50 bases upstream of the cleavage site for genes expressed in both *bam*^-/-^; *hs-Bam-HA* flies without heat shock and 48h PHS that had identical 3’ cleavage site choice for the predominant cleavage site in all biological replicates between *bam* and 48h PHS 3’Seq libraries. A 2^nd^ order Markov model was used as the background model.

### Polysome Fractionation

For each biological replicate of the polysome fractionation, approximately 1,200 testes (dissected from 600 flies) were used, combining 10 tubes of 120 testes (flash-frozen in liquid nitrogen and stored at -80°C) during the lysis step. Tissue was lysed in buffer containing 25 mM Tris pH 7.5, 15 mM MgCl2, 150 mM NaCl, 1% Triton X-100, 1 mM DTT, 8% Glycerol, and 0.2 mg/mL Cycloheximide at 4°C using a 27 gauge 1-1/2^”^ syringe to disrupt the tissue and then the lysate was incubated for 30 mins at 4°C while rocking. 10% of the supernatant was saved for input and the remaining supernatant was loaded onto a 50mL sucrose gradient (10%-40%). Samples were spun down in an ultracentrifuge at 40,000 rpm for 2.5 hrs at 4°C. Sequential 10 drop fractions were collected from the top using a Brandel gradient fractionator, with absorbance at A260 nm measured continuously during collection to assess RNA concentration. RNA was isolated by acid phenol-chloroform extraction and fractions were combined based on the absorbance profiles at A260 nm to create 6 groups: free, 40S, 60S, 80S, 2-3 ribosomes and 4+ ribosomes. 3’ Seq libraries were created using the QuantSeq FWD kit (Lexogen) and reads were mapped and analyzed with the same pipeline outlined in ‘3’ Seq Data Analysis’ (Figure S1A). Sequencing depth was sufficient for analysis of all biological replicates of 24h, 48h and 72h PHS (Figure S6A-C) except for one replicate of the 72h PHS 60S polysome fraction, which was discarded due to the low number of reads mapping to gene 3’UTRs. Read distributions are shown for the orb locus in the polysome profiles from 24h, 48h and 72h PHS in Figure S6D-F.

### Polysome fractionation analysis

3’ Seq reads were normalized to the read count in each library and then normalized across the polysome profile to analyze all transcripts independent of the overall expression. Hierarchical clustering was used to group transcripts with similar polysome profiles. Clusters were then rearranged to reflect the order of the biological process (i.e., the cluster containing transcripts enriched in the free fraction was ordered above the cluster containing transcripts enriched in the 40S fraction, etc.).

### Alternative 3’ cleavage analysis using Fly Cell Atlas (FCA) 10X data

Since the FCA 10X reads were always shifted approximately 200 bases upstream of the CSs identified by 3’ Seq (Figure S3A and S3B), we summed all reads, up to 250 bases upstream of the proximal and distal CSs, to reflect the expression of the proximal and distal CSs in the FCA data set. We then calculated the ratio of proximal:distal CS usage using all reads from nuclei in cluster 25 of Leiden 6.0 (corresponding to spermatogonia) and all reads from nuclei in cluster 35 of Leidein 6.0 (corresponding to early to mid-stage spermatocytes).

### Immunofluorescence staining

For whole-mount staining, testes from 0-1 day old males were dissected in 1X phosphate-buffered saline (PBS) and incubated with 4% formaldehyde for 20 minutes at room temperature. After fixation, the testes were washed once in PBST (PBS with 0.1% Triton X-100) and permeabilized by incubation with PBS with 0.3% Triton X-100 and 0.6% sodium deoxycholate for 30 minutes at room temperature. After permeabilization, testes were washed once in PBST and blocked for 30 minutes with PBST containing 3% bovine serum albumin (BSA), then incubated overnight at 4°C with desired primary antibodies in PBST with 3% BSA. After overnight incubation in primary antibody, testes were washed three times with PBST, incubated with secondary antibodies conjugated with Alexa fluorophores (Alexa Fluor-488, -568, -647 from Molecular Probes) for 2 hours at room temperature in the dark while rocking, then washed three times in PBST and mounted on glass slides with mounting medium containing DAPI (VECTASHIELD, Vector Labs, Cat# H-1200).

The sources and dilutions of primary antibodies used were as follows: Vasa (goat, 1:100; Santa Cruz Biotechnology, Cat# dc-13); Lola-F (mouse, 1:100, 1F1-1D5, DSHB); Kumgang (rabbit, 1;400) (Kim et al. 2017); NudE (rabbit, 1:200) (Wainman et al. 2009); Orb (mouse, orb 4H8 and orb 6H4, DSHB, 1:30), Numb (guinea-pig, 1:500) (O’Connor-Giles and Skeath 2003).

### Generation of transgenic reporters

The sequence encoding destabilized GFP (Li et al. 1998) flanked with EcoRI and NotI restriction sites were synthesized. Additionally, the sequence encoded in the endogenous *dco* 3’UTR (total of 1857 nt, “*dco* 3’UTR WT”) flanked with Not1 and Kpn1 restriction sizes and a second similar sequence with one base pair change, the 113^th^ T in the *dco* 3’UTR into an A (“*dco* 3’UTR can PAS*”), were synthesized. To create the expression constructs, the pUASTattb vector (Bischof et al. 2007) was digested with EcoRI and KpnI enzymes and gel purified, then ligated with the sequences encoding the destabilized GFP and one or the other of the *dco* 3’UTR sequences. These constructs were injected into fly embryos carrying the attp40 site by BestGene and fly lines that had stably introduced these transgenes into the germline were retained.

### Fluorescence microscopy and image analysis

Images of fixed and immunostained testes were taken with either a Leica SP2 or SP8 confocal microscope. Images of native GFP and mRFP fluorescence in unfixed testes were taken with a Leica SP8 microscope. Laser intensity and detector gain for the confocal microscope were adjusted for each experiment to ensure that the signal was in the linear range and not saturated. Once image acquisition settings were determined, the same settings were maintained for the entire set of images to be compared. Immunofluorescence images were processed using ImageJ with all samples to be compared going through the same processing.

For live imaging of testes expressing fluorescent reporters, newly dissected testes in 1X PBS were immediately placed on a coverslip. To prevent squashing during imaging, two clean coverslips were securely taped on a slide with a gap left in between and the coverslip with the dissected testes was placed on the slide in the gap with edges overlapping the taped coverslips. Testes were imaged on a Leica SP8 confocal, taking approximately 40 z-slices with a separation of 2 microns between each, focused on the tip of the testis. Quantification of fluorescence was performed using the program Volocity (PerkinElmer) to sum fluorescence of a 120 μm x 120 μm area positioned to cover the tip of the testis across z-stacks. GFP reporter fluorescence was normalized to His2Av-mRFP fluorescence in each sample.

### qRT-PCR and 3’RACE

RNA was extracted from ∼30 pairs of testes by phenol-chloroform separation using Trizol (Life Technologies, Cat# 15596-026). 0.5μg of total RNA was used to make cDNA by using Ready-To-Go You Prime First-Strand Beads (GE Healthcare, Cat# 27-9264-01) with 0.1 μg of oligo dT or 3’RACE adapter per reaction (Invitrogen FirstChoice® RLM-RACE Kit, Cat#AM1700). After reverse transcription, cDNA was diluted 1/100 in filtered water and 5-10 μL of diluted cDNA was used per qPCR reaction. To perform qRT-PCR, we mixed 2X SensiFAST SYBR Hi-ROX reagents (Bioline, Cat # BIO-92020) with cDNA and a final concentration of 125nM for each primer and measured fluorescence using the 7500 Fast Real-Time PCR System from Applied Biosystems. The relative abundance of the transcripts was calculated by the ΔΔCt method. In all qRT-PCR experiments, error bars represent the standard deviation from the mean in three independent biological replicates.

For 3’ RACE, cDNA was generated following manufacturer instructions (Invitrogen FirstChoice® RLM-RACE Kit). Briefly, first strand synthesis reaction was carried out with the 3’ RACE adapter (5’-GCGAGCACAGAATTAATACGACTCACTATAGGT_12_VN-3’) at 42°C for one hour. The outer PCR reaction was carried out with an outer *lolaF* isoform-specific primer (5’-CGATGACGAAGACGAGACATT-3’) and the outer primer (5’-GCGAGCACAGAATTAATACGACT-3’) for 35 cycles. 1/20 of the PCR product was used in the subsequent inner PCR reaction with an inner *lolaF* isoform-specific primer (5’-GATGAACGAGGAGTGGAACAT-3’) as well as the inner primer (5’-CGCGGATCCGAATTAATACGACTCACTATAGG-3’) for 35 cycles.

### Protein-protein interaction network and Gene Ontology (GO) Analysis

Protein-protein interaction network analysis was conducted by using STRING (version 11.5) (Szklarczyk et al. 2019; Szklarczyk et al. 2021) using a minimum required interaction score of 0.900 (highest confidence) or 0.700 (high confidence). Active interaction sources included in the interaction analysis were: (i) experimentally determined interactions, (ii) curated pathway databases and (iii) co-expression evidence. GO term enrichments were determined using STRING. As input, the 531 transcripts that undergo APA to produce transcripts with shorter 3’ UTRs in differentiating spermatocytes or 200 transcripts that shorten their 3’UTR from 24h PHS to 48h PHS in the time-course and their translation activity goes from ON to OFF were used.

## Acknowledgements

We thank Dr. Maurizio Gatti for antibodies against NudE, Dr. James Skeath for antibodies against Numb and Drs. Kristen and Jørgen Johansen for antibodies against LolaF. We thank Dr. Maria Barna for suggesting the polysome fractionation approach, Dr. Kathrin Leppek, Dr. Gerald Tiu, and Dr. Shifeng Xue in the Barna laboratory for technical assistance with polysome fractionation, and Dr. Ariel Bazzini for advice on interpreting effects of miRNAs on polysome distribution of their mRNA targets. We acknowledge Dr. Erin Davies and William Joo for their initial observations on alternative *lolaF* isoforms and LolaF protein expression in *Drosophila* testes and Dr. Erin Sanders for her contributions to analysis of Numb protein expression in testes during her rotation. We are grateful to the Bloomington *Drosophila* Stock Center and the Vienna *Drosophila* RNAi Center for fly stocks, the Developmental Studies Hybridoma Bank for several antibodies, Cheryl Smith from Arendt Sidow’s lab for insightful help on library preparation, Reuven Agami for advice on library preparation and 3’Seq analysis, Lars Steinmetz for comments, and members of the Fuller laboratory for helpful discussions and input on the manuscript. CWB was supported by a Stanford Graduate Fellowship and NIH T32AR007422 (PI: Dr. Paul Khavari). GHO was supported by a Latin American Pew Fellowship, an American Heart Association (AHA) postdoctoral fellowship and Becas Chile (ANID). LG was supported by an American-Italian Cancer Foundation Post-Doctoral Research fellowship, year 2021-2022. This work was supported by NIH grants R21HD079970, 1R01GM124054, R35GM136433 and funds from the Reed-Hodgson Professorship in Human Biology to MTF.

## Author Contributions

C.W.B., G.O.H., and L.G. performed experiments. C.W.B. and G.O.H. analyzed data. C.W.B. performed all 3’ Seq and polysome profiling experiments. C.W.B. performed all the bioinformatic analysis. G.R. created initial pipeline for 3’Seq analysis with supervision from J.B.L. G.O.H. performed 3’RACE experiments. C.W.B. and G.O.H. designed and analyzed 3’UTR reports. L.G. performed Immunofluorescence experiments. G.O.H. performed functional interaction network analysis. C.W.B. and G.O.H. prepared the figures. C.W.B., G.O.H. and M.T.F. designed the study. C.W.B., G.O.H., and M.T.F. wrote the manuscript, together with feedback from A.G., P.O. and J.B.L. The manuscript was reviewed by all authors.

