## Supplemental Files and Figures for "Developmentally regulated alternate 3’ end cleavage of nascent transcripts controls dynamic changes in protein expression in an adult stem cell lineage"

A

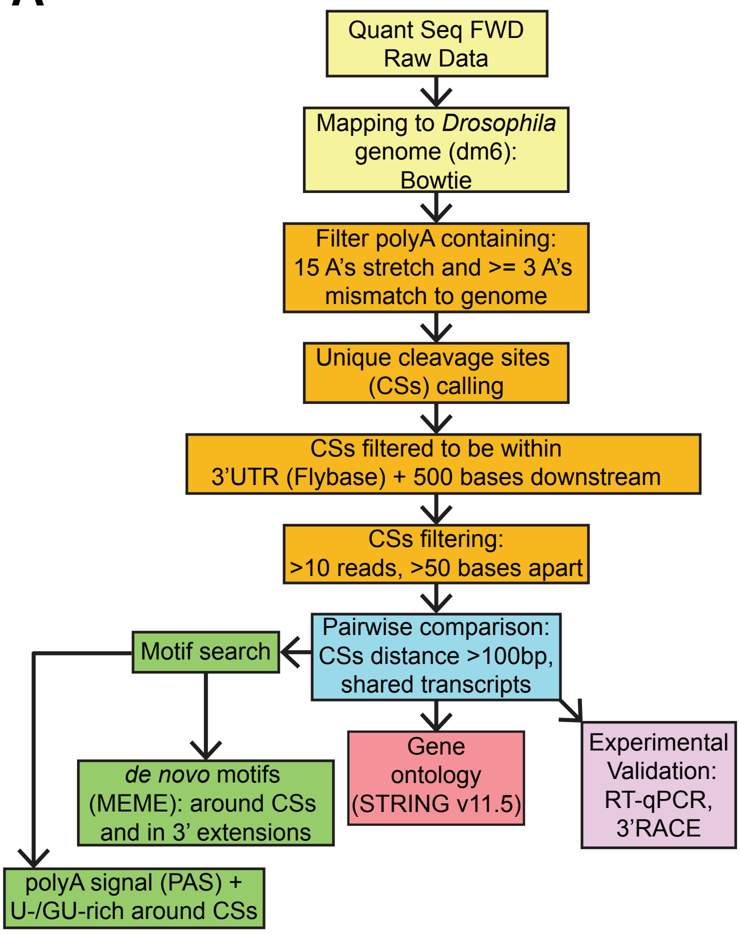


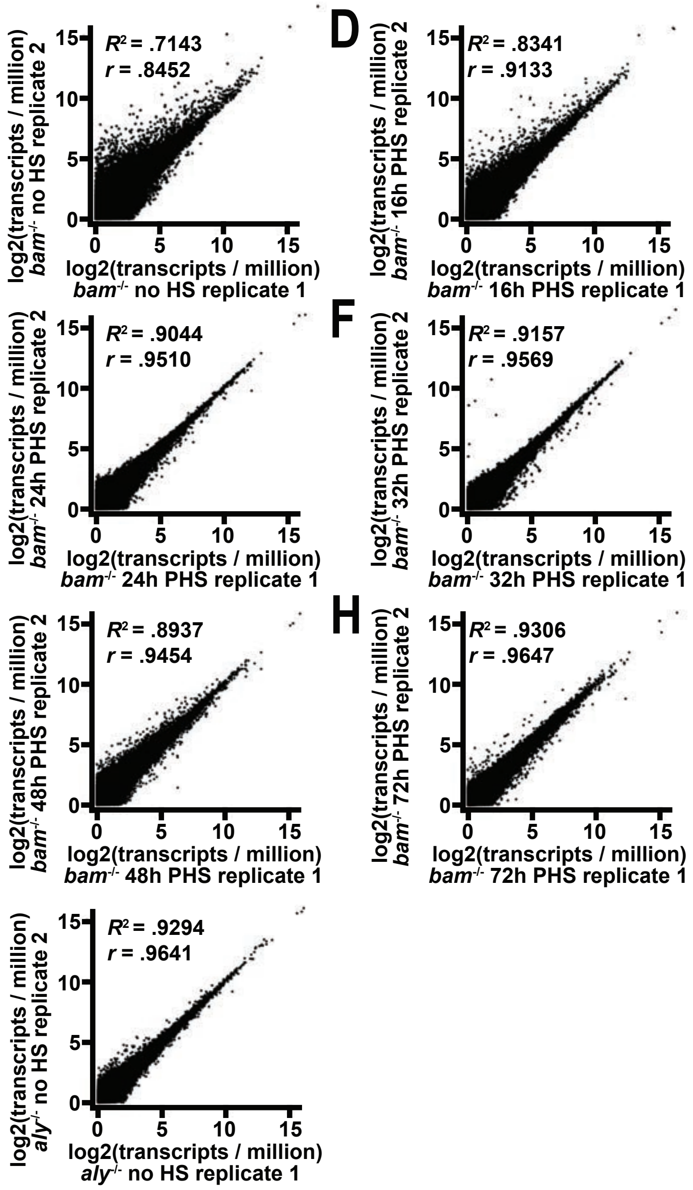


B

| 3'Seq Library Genotype | Uniquely mapped reads | polyA-spanning reads | Unique cleavage sites (CSs) | Filtered CSs (>10 reads, >50 bases apart, 3'UTR +500bp) | shared | Number of genes corresp. to filtered CSs | shared |
|---|---|---|---|---|---|---|---|
| *bam*$^{-/-}$;hs>Bam - no HS R1 | 57,876,832 | 33,032,825 | 305,192 | 20,197 | 12,613 | 9,127 | 8,516 |
| *bam*$^{-/-}$;hs>Bam - no HS R2 | 63,573,917 | 29,426,622 | 275,282 | 20,367 | | 9,328 | |
| *bam*$^{-/-}$;hs>Bam - 16h PHS R1 | 70,867,617 | 34,696,012 | 234,925 | 21,965 | 13,586 | 9,578 | 9,081 |
| *bam*$^{-/-}$;hs>Bam - 16h PHS R2 | 76,929,724 | 32,232,886 | 326,909 | 23,248 | | 10,341 | |
| *bam*$^{-/-}$;hs>Bam - 24h PHS R1 | 76,322,920 | 32,063,864 | 277,788 | 21,968 | 14,318 | 9,572 | 9,107 |
| *bam*$^{-/-}$;hs>Bam - 24h PHS R2 | 77,676,349 | 34,650,023 | 301,098 | 22,912 | | 9,749 | |
| *bam*$^{-/-}$;hs>Bam - 32h PHS R1 | 75,036,927 | 34,084,988 | 387,638 | 22,819 | 15,196 | 9,944 | 9,287 |
| *bam*$^{-/-}$;hs>Bam - 32h PHS R2 | 55,562,813 | 25,583,523 | 295,071 | 21,714 | | 9,881 | |
| *bam*$^{-/-}$;hs>Bam - 48h PHS R1 | 59,579,236 | 29,114,345 | 384,518 | 24,570 | 18,008 | 11,360 | 11,020 |
| *bam*$^{-/-}$;hs>Bam - 48h PHS R2 | 95,752,424 | 46,832,920 | 579,143 | 27,444 | | 11,765 | |
| *bam*$^{-/-}$;hs>Bam - 72h PHS R1 | 81,691,614 | 38,859,460 | 556,399 | 24,805 | 18,904 | 11,505 | 11,020 |
| *bam*$^{-/-}$;hs>Bam - 72h PHS R2 | 68,508,829 | 32,992,852 | 635,937 | 23,437 | | 11,303 | |
| *aly*$^{-/-}$ - no HS R1 | 94,544,416 | 43,647,401 | 553,400 | 24,796 | 16,217 | 10,559 | 9,968 |
| *aly*$^{-/-}$ - no HS R2 | 86,107,077 | 40,792,827 | 561,278 | 24,449 | | 10,564 | |

J

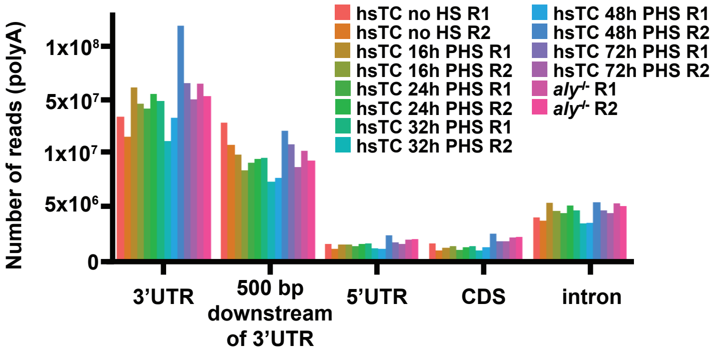


K

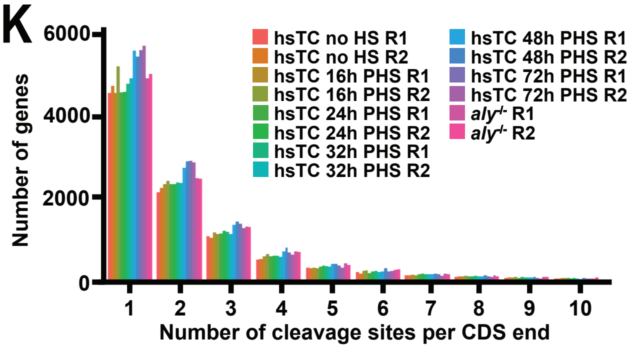


Figure SOM1.

**Supplementary Materials – Legends**

**Figure S1. Pipeline for 3' Seq library preparation and analysis of 3' cleavage sites obtained from 3' Seq**. (A) 3' Seq bioinformatic analysis pipeline used in this paper, preceded by the synthesis of cDNA and library prep using the QuantSeq FWD kit and sequencing on the NextSeq Illumina platform. (B) Quantification of numbers of reads, cleavage sites (CSs) and corresponding genes identified using the above bioinformatic analysis pipeline from two biological replicates (indicated as R1 and R2) of libraries prepared from testes of indicated genotypes/heat shock time course (hsTC) conditions. (C-I) Scatterplot of expression levels for all transcripts expressed with at least 1 transcript per million in both biological replicates (R1 and R2) of indicated genotypes/hsTC treatment conditions. ($R^2$) Coefficient of determination. (*r)* Pearson correlation coefficient. (J) Location of 3' end cleavage sites in indicated genotypes/hsTC treatment conditions with respect to gene structure (i.e., in 3'UTR, up to 500 bases downstream of the 3'UTR, 5'UTR, CDS and introns) in both biological replicates (R1 and R2). (K) Number of 3' end cleavage sites associated with unique 3' CDS ends in both biological replicates (R1 and R2) of indicated genotypes/hsTC treatment conditions.

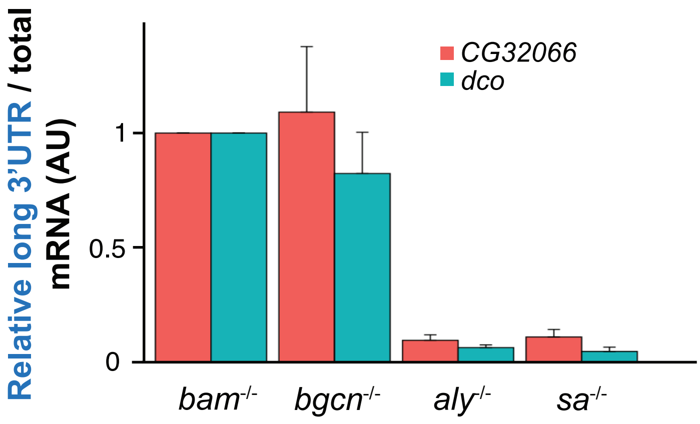
Relative long 3'UTR / total mRNA (AU)
CG32066
dco
1
0.5
0
bam-/-
bgcn-/-
aly-/-
sa-/-

Figure SOM2.

**Figure S2. Validation of mRNA isoforms with shorter 3'UTRs in spermatocytes vs. spermatogonia by qRT-PCR.** (A) RT-PCR ratio of long 3'UTR isoform relative to total transcript for *CG32066* (red) and *dco* (cyan) in the indicated mutant testes. *bam* and *bgcn* mutant testes are both highly enriched for proliferating spermatogonia and lack spermatocytes. *aly* and *sa* mutant testes are enriched for mature spermatocytes, contain some spermatogonia, and lack spermatids. Relative long/total ratio expressed as mean + standard deviation and long/total ratio in *bam* mutant set to 1.

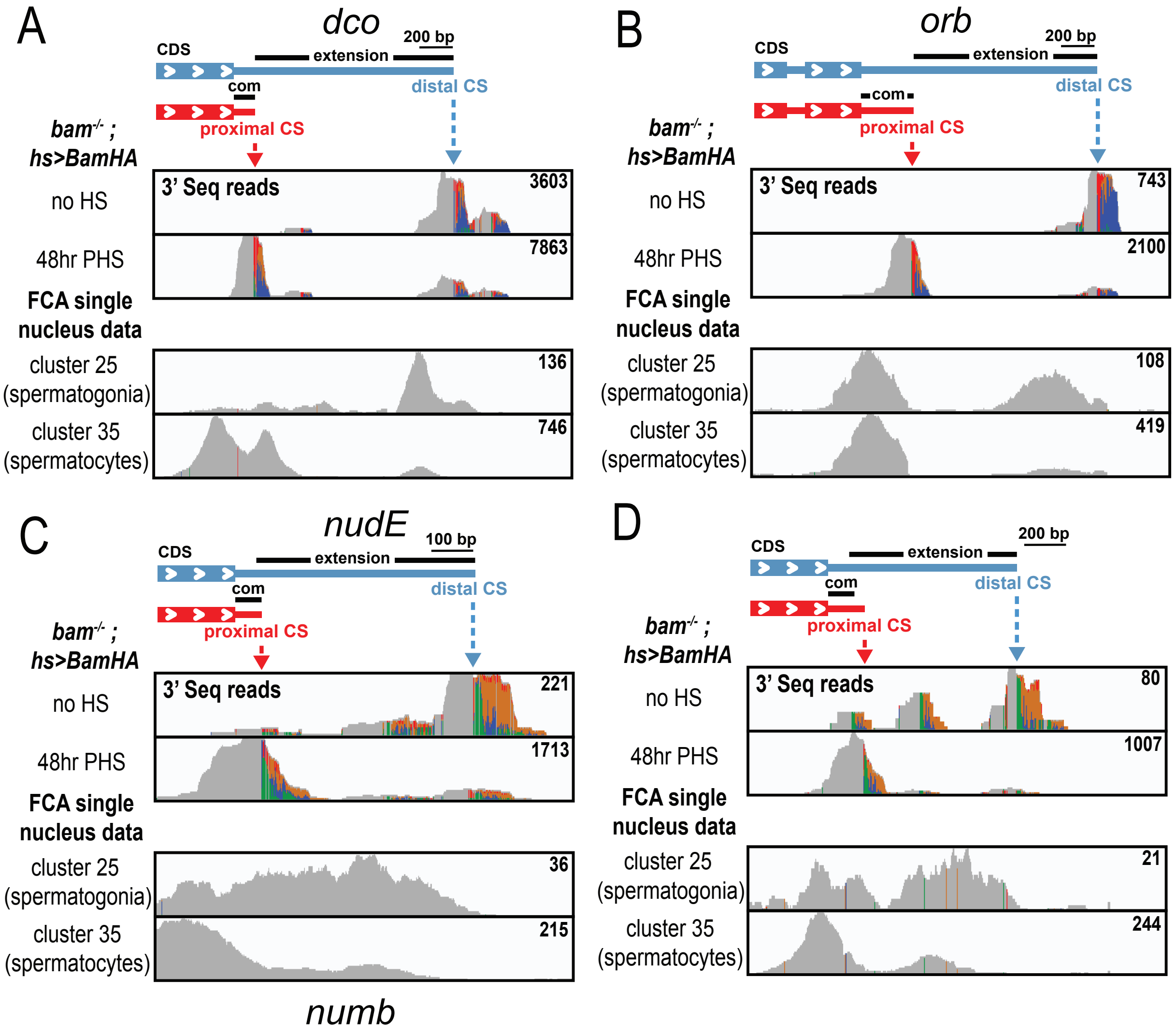


E

Global Analysis of FCA reads from
cluster 35 (spermatocytes)
vs. cluster 25 (spermatogonia)

- Lower ratio of proximal:distal CS usage in cluster 35 vs 25 (FCA)
- Higher ratio of proximal:distal CS usage in cluster 35 vs 25 (FCA)

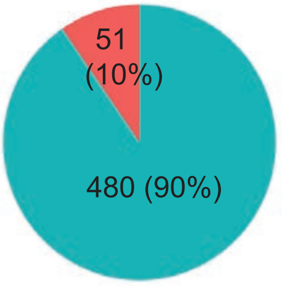


$$= \frac{\text{(proximal CS read count in cluster 35) / (distal CS read count in cluster 35)}}{\text{(proximal CS read count in cluster 25) / (distal CS read count in cluster 25)}}$$

Figure SOM3.

**Figure S3. Analysis of Fly Cell Atlas snRNA-Seq data supports alternative 3' cleavage in spermatocytes compared to spermatogonia.** (A, B, C, D) 3' Seq tracks from one of two biological replicates from testes from *bam* hsTC flies at indicated times PHS, as well as tracks from Fly Cell Atlas (FCA) single nucleus sequencing data plotted on the 3' genomic regions for (A) *dco*, (B) *orb*, (C) *nudE*, and (D) *numb*. On top, a gene model scheme representing the two most abundant isoforms of 3'UTR for each gene. Proximal (red arrowhead) and distal (blue arrowhead) cleavage sites (CS) were predicted from the most highly used cleavage site in $bam^{\Delta 86/1}$*; hs-Bam* testes without heat shock and 48hr PHS, respectively. (E) Pie chart indicating the fraction of the 531 genes identified in this study as undergoing alternative 3' cleavage in the hsTC that show a higher ratio of proximal to distal CS expression in the FCA single nucleus sequencing data in spermatocytes (cluster 35, Leiden 6.0) vs spermatogonia (cluster 25, Leiden 6.0).

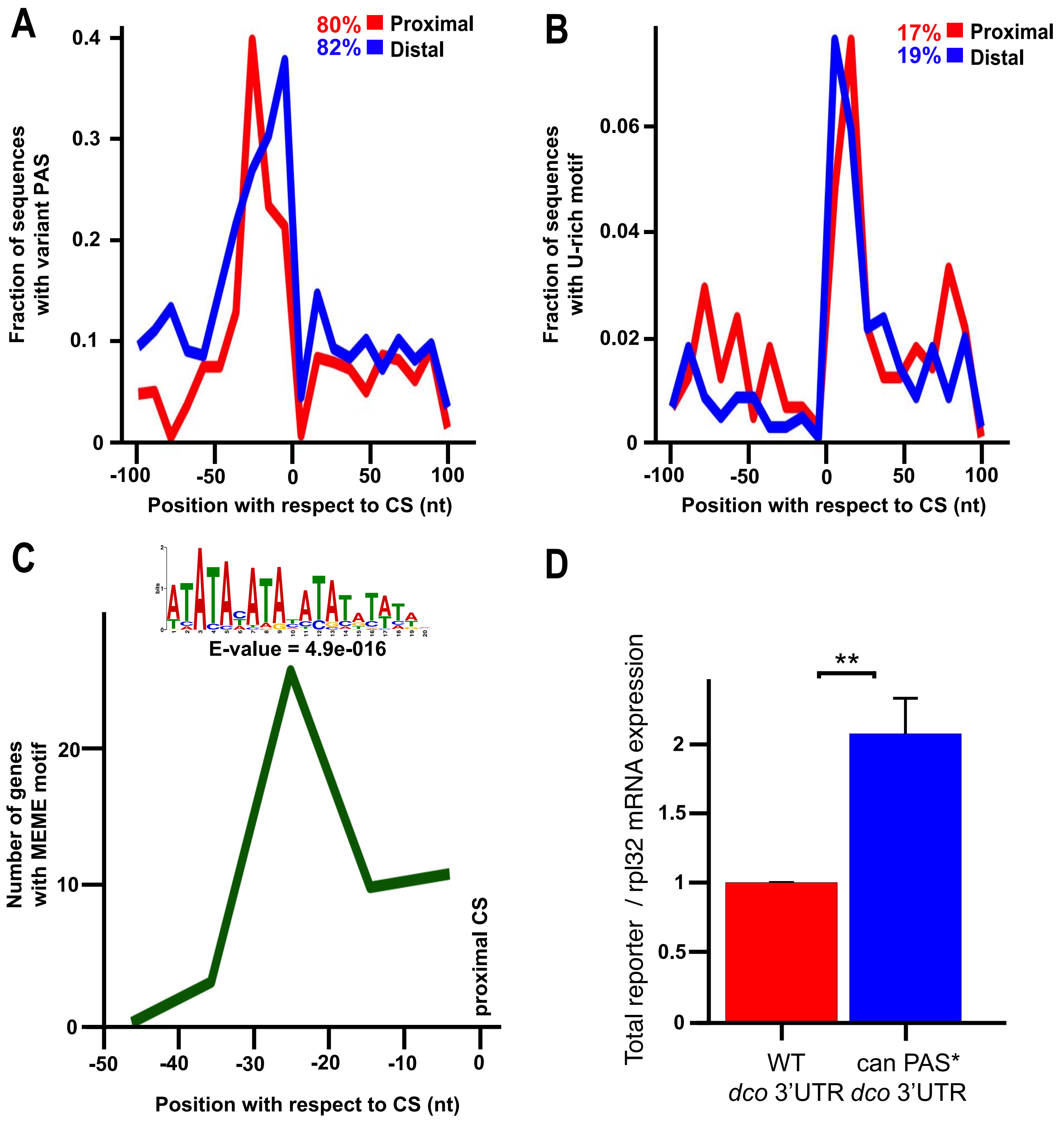
A
80% Proximal
82% Distal
Fraction of sequences with variant PAS
0.4
0.3
0.2
0.1
0
-100
-50
0
50
100
Position with respect to CS (nt)
B
17% Proximal
19% Distal
Fraction of sequences with U-rich motif
0.06
0.04
0.02
0
-100
-50
0
50
100
Position with respect to CS (nt)
C
E-value = 4.9e-016
Number of genes with MEME motif
20
10
0
proximal CS
-50
-40
-30
-20
-10
0
Position with respect to CS (nt)
D
**
Total reporter / rpl32 mRNA expression
2
1.5
1
0.5
0
WT
dco 3'UTR
can PAS*
dco 3'UTR

Figure SOM4

**Figure S4. Motif analysis of sequences around the proximal and distal cleavage sites of genes that undergo stage-specific APA.** (A) Presence and position of variant PAS motifs (ATTAAA, TATAAA, AGTAAA, TTTAAA, AATATA, CATAAA, AATACA, GATAAA, AAGAAA, AAAAAG, AAAACA, AATAGA) 100 nt upstream of proximal (red) and distal (blue) cleavage sites (CSs). Percentages indicate the number of transcripts with a variant PAS between 10 - 40 nt upstream of the indicated CS. (B) Presence and position of U-rich sequences 100 nt upstream and downstream of proximal (in red) and distal (blue) CSs. Percentages indicate the number of transcripts with U-rich sequences between 0 - 50 nt downstream of the indicated CS. (C) Location and identity of the most prevalent motif identified by MEME analysis of the 50 nt region upstream of the proximal 3' end cleavage site of genes that undergo APA relative to the 50 nt region upstream of the cleavage site of the set of genes that do not undergo APA but are expressed in *bam*$^{-/-}$*; hs-Bam* flies without heat shock and 48h PHS. (D) RT-PCR ratio of total mRNA produced in testes from indicated reporters expressed under control of *nos-Gal4* relative to housekeeping *rpl32* mRNA. Statistical significance determined by two-tailed student's t-test (** indicates $p$-value < 0.01). Relative reporter/*rpl32* ratio expressed as mean + standard deviation (3 independent biological replicates) and reporter/*rpl32* ratio in WT *dco* 3'UTR reporter was set to 1.

**A**

| 3'Seq Library Genotype and Polysome fraction | Uniquely mapped reads | polyA-spanning reads | Unique cleavage sites (CSs) |
|---|---|---|---|
| 24h PHS - free R1 | 38,538,648 | 21,160,206 | 289,232 |
| 24h PHS - free R2 | 41,292,474 | 22,389,941 | 258,755 |
| 24h PHS - 40S R1 | 38,834,866 | 18,137,694 | 167,663 |
| 24h PHS - 40S R2 | 43,673,370 | 19,851,298 | 191,002 |
| 24h PHS - 60S R1 | 33,731,842 | 15,586,366 | 105,227 |
| 24h PHS - 60S R2 | 51,556,187 | 25,488,408 | 131,813 |
| 24h PHS - 80S R1 | 41,500,610 | 19,258,202 | 173,793 |
| 24h PHS - 80S R2 | 41,821,095 | 18,968,231 | 206,032 |
| 24h PHS - 2-3 ribo. R1 | 39,186,134 | 18,016,964 | 188,847 |
| 24h PHS - 2-3 ribo. R2 | 40,554,828 | 17,008,582 | 158,204 |
| 24h PHS - 4 plus ribo. R1 | 39,475,154 | 16,422,624 | 140,857 |
| 24h PHS - 4 plus ribo. R2 | 36,249,478 | 15,187,730 | 140,045 |

**B**

| 3'Seq Library Genotype and Polysome fraction | Uniquely mapped reads | polyA-spanning reads | Unique cleavage sites (CSs) |
|---|---|---|---|
| 48h PHS - free R1 | 36,178,482 | 18,043,456 | 487,506 |
| 48h PHS - free R2 | 32,516,531 | 16,064,486 | 288,451 |
| 48h PHS - 40S R1 | 30,928,418 | 15,585,328 | 366,273 |
| 48h PHS - 40S R2 | 33,209,713 | 15,499,165 | 270,265 |
| 48h PHS - 60S R1 | 26,444,953 | 11,330,781 | 163,337 |
| 48h PHS - 60S R2 | 33,865,303 | 15,249,998 | 158,341 |
| 48h PHS - 80S R1 | 33,040,404 | 14,687,626 | 243,850 |
| 48h PHS - 80S R2 | 33,968,717 | 14,029,076 | 180,044 |
| 48h PHS - 2-3 ribo. R1 | 47,345,431 | 21,005,490 | 229,957 |
| 48h PHS - 2-3 ribo. R2 | 34,543,819 | 13,684,553 | 163,366 |
| 48h PHS - 4 plus ribo. R1 | 31,909,184 | 14,857,352 | 223,427 |
| 48h PHS - 4 plus ribo. R2 | 36,433,448 | 14,922,218 | 172,796 |

**C**

| 3'Seq Library Genotype and Polysome fraction | Uniquely mapped reads | polyA-spanning reads | Unique cleavage sites (CSs) |
|---|---|---|---|
| 72h PHS - free R1 | 47,250,464 | 23,853,637 | 579,243 |
| 72h PHS - free R2 | 34,301,459 | 16,327,687 | 465,263 |
| 72h PHS - 40S R1 | 40,807,017 | 18,646,323 | 329,665 |
| 72h PHS - 40S R2 | 43,714,037 | 20,113,412 | 194,351 |
| 72h PHS - 60S R1 | 39,517,547 | 19,187,504 | 123,041 |
| 72h PHS - 60S R2 | 42,925,418 | 17,591,114 | 66,884* |
| 72h PHS - 80S R1 | 43,333,116 | 15,444,557 | 288,765 |
| 72h PHS - 80S R2 | 39,119,098 | 15,444,557 | 265,717 |
| 72h PHS - 2-3 ribo. R1 | 31,397,775 | 13,124,985 | 278,088 |
| 72h PHS - 2-3 ribo. R2 | 35,602,394 | 11,566,218 | 187,243 |
| 72h PHS - 4 plus ribo. R1 | 30,785,197 | 13,053,383 | 335,177 |
| 72h PHS - 4 plus ribo. R2 | 41,336,992 | 15,947,125 | 375,909 |

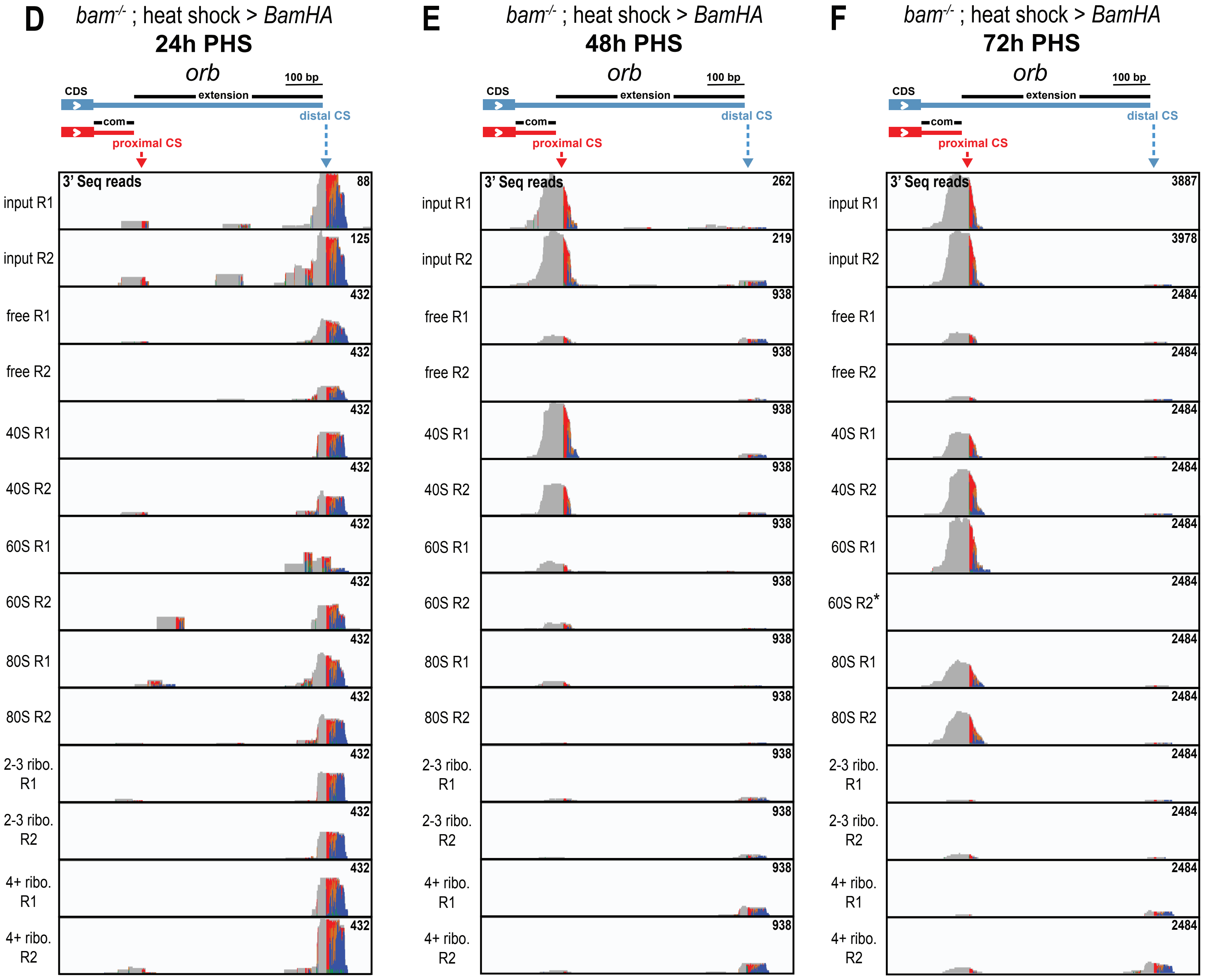


Figure SOM5.

**Figure S5. 3' Seq following polysome fractionation.** (A-C) Quantification of the number of reads, cleavage sites (CSs) and corresponding genes identified using bioinformatic analysis pipeline from Figure S1A. 3' Seq data from two biological replicates (R1 and R2) of libraries prepared from indicated genotypes/hs-TC treatments. (D-F) 3' Seq tracks plotted on the 3' genomic region of *orb* for two biological replicates (R1 and R2) of the polysome fractions indicated from testes from *bam* hsTC flies dissected at the indicated times PHS. Successive polysome fractions are displayed from the top (free RNA) to the bottom (polysomal fractions with 4 or more ribosomes). Asterisk indicates one biological replicate of the 3' Seq 60S fraction from 72h PHS that was discarded due to the low number of genes detected.

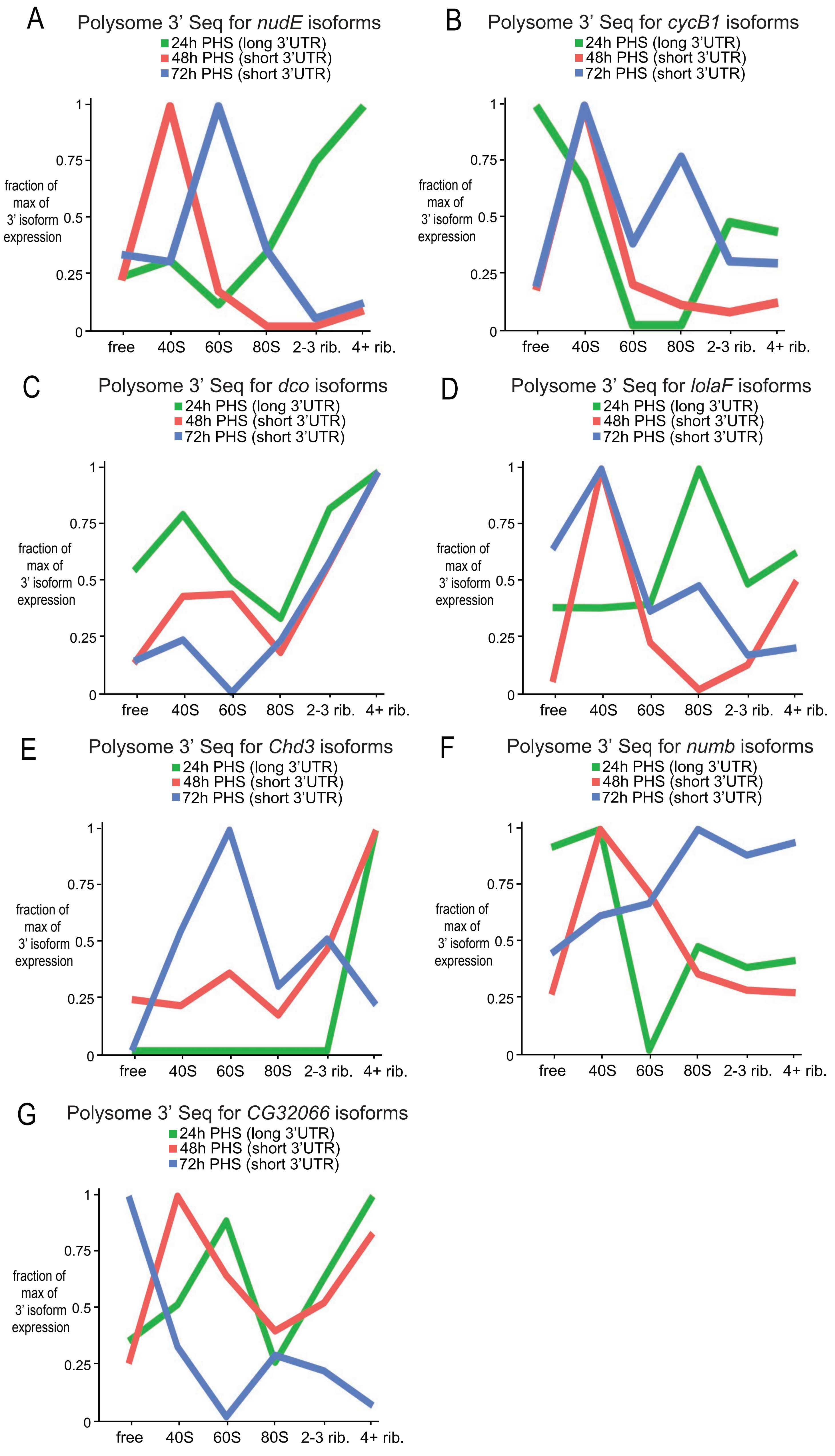
A
Polysome 3' Seq for nudE isoforms
24h PHS (long 3'UTR)
48h PHS (short 3'UTR)
72h PHS (short 3'UTR)
fraction of max of 3' isoform expression
0
0.25
0.5
0.75
1
free
40S
60S
80S
2-3 rib.
4+ rib.
B
Polysome 3' Seq for cycB1 isoforms
24h PHS (long 3'UTR)
48h PHS (short 3'UTR)
72h PHS (short 3'UTR)
fraction of max of 3' isoform expression
0
0.25
0.5
0.75
1
free
40S
60S
80S
2-3 rib.
4+ rib.
C
Polysome 3' Seq for dco isoforms
24h PHS (long 3'UTR)
48h PHS (short 3'UTR)
72h PHS (short 3'UTR)
fraction of max of 3' isoform expression
0
0.25
0.5
0.75
1
free
40S
60S
80S
2-3 rib.
4+ rib.
D
Polysome 3' Seq for lolaF isoforms
24h PHS (long 3'UTR)
48h PHS (short 3'UTR)
72h PHS (short 3'UTR)
fraction of max of 3' isoform expression
0
0.25
0.5
0.75
1
free
40S
60S
80S
2-3 rib.
4+ rib.
E
Polysome 3' Seq for Chd3 isoforms
24h PHS (long 3'UTR)
48h PHS (short 3'UTR)
72h PHS (short 3'UTR)
fraction of max of 3' isoform expression
0
0.25
0.5
0.75
1
free
40S
60S
80S
2-3 rib.
4+ rib.
F
Polysome 3' Seq for numb isoforms
24h PHS (long 3'UTR)
48h PHS (short 3'UTR)
72h PHS (short 3'UTR)
fraction of max of 3' isoform expression
0
0.25
0.5
0.75
1
free
40S
60S
80S
2-3 rib.
4+ rib.
G
Polysome 3' Seq for CG32066 isoforms
24h PHS (long 3'UTR)
48h PHS (short 3'UTR)
72h PHS (short 3'UTR)
fraction of max of 3' isoform expression
0
0.25
0.5
0.75
1
free
40S
60S
80S
2-3 rib.
4+ rib.

Figure SOM6.

**Figure S6. Polysome profiles of genes that undergo APA.** (A-G) Line graphs of the distribution of the long 3'UTR isoform (green, 24h PHS) and short 3'UTR isoform (red, 48h PHS; blue, 72h PHS) in the indicated polysome fractions for (A) *nudE*, (B) *cycB1*, (C) *dco*, (D) *lola-F*, (E) *Chd3*, (F) *numb* and (G) *CG32066*.

A

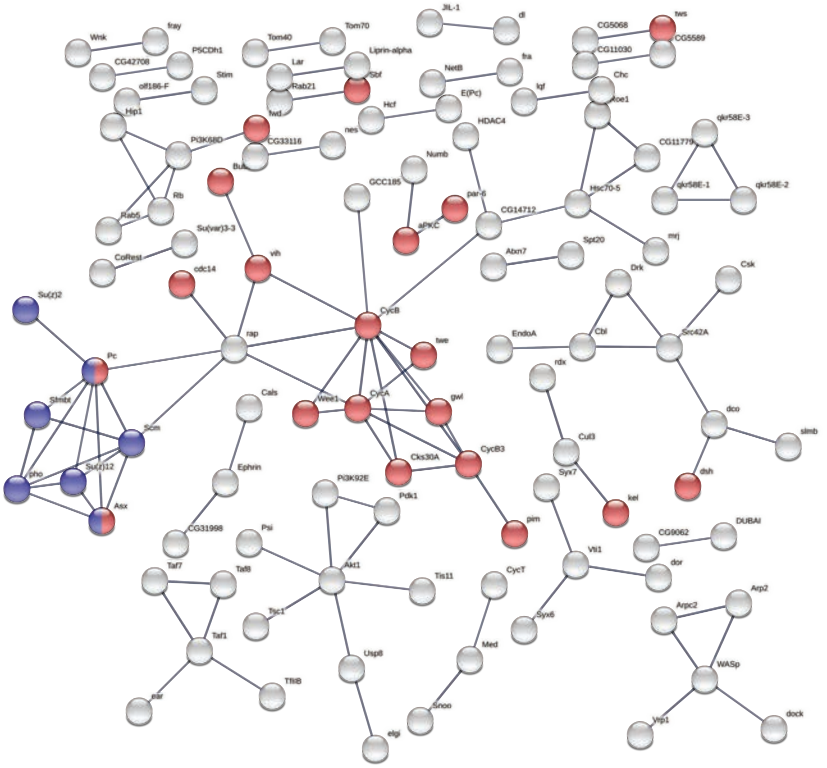

PPI enrichment *p*-value: 2.02e-10

B

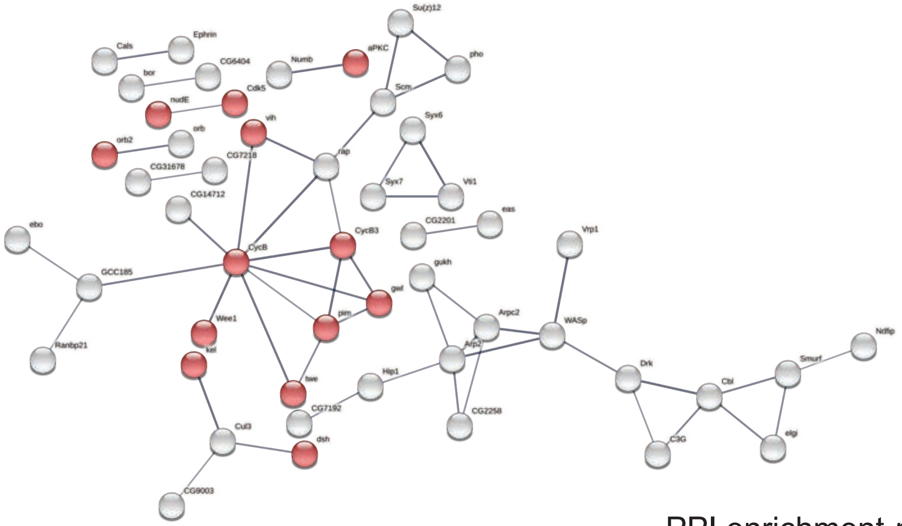

PPI enrichment *p*-value: 2.46e-11

Figure SOM7.

**Figure S7. Functional protein association network of genes that shorten their 3'UTR during germline differentiation in the hsTC.** (A) Protein-protein interaction network of the 531 genes that undergo APA resulting in shorter 3'UTRs as they progress in the heat shock time course (hsTC) conditions. The network displays the predictions of protein-protein and other experimentally determined interactions, including predictions from curated pathway databases and co-expression modules. Highlighted in red are the nodes that belong to the "Cell cycle" Gene Ontology Biological Process category (GO:0007049), FDR: 3.12e-07. Highlighted in blue are the nodes that belong to the "PcG protein complex" Gene Ontology Cellular Component category (GO:0031519), FDR: 7.6e-03. (B) Functional protein association network of genes that shorten their 3'UTR from 24h PHS to 48h PHS in the time-course and their translation activity goes from ON to OFF. Highlighted in red are the nodes that belong to the "Cell cycle process" Gene Ontology Biological Process category (GO:0022402), FDR: 4.5e-04. STRING database (version 11.5) (Szklarczyk et al. 2021) was used for this analysis with the minimum required interaction score of (A) 0.900 (highest confidence) and (B) of 0.700 (high confidence). Line thickness indicates the strength of data support. Disconnected nodes in the network are not shown.
